# Directed evolution of the multicopper oxidase laccase for cell surface proximity labeling and electron microscopy

**DOI:** 10.1101/2024.10.29.620861

**Authors:** Song-Yi Lee, Heegwang Roh, David Gonzalez-Perez, Mason R. Mackey, Keun-Young Kim, Daniel Hoces, Colleen N. McLaughlin, Stephen R. Adams, Khanh Nguyen, David J. Luginbuhl, Liqun Luo, Namrata D. Udeshi, Steven A. Carr, Rogelio A. Hernández-López, Mark H. Ellisman, Miguel Alcalde, Alice Y. Ting

## Abstract

Enzymes that oxidize aromatic substrates have shown utility in a range of cell-based technologies including live cell proximity labeling (PL) and electron microscopy (EM), but are associated with drawbacks such as the need for toxic H_2_O_2_. Here, we explore laccases as a novel enzyme class for PL and EM in mammalian cells. LaccID, generated via 11 rounds of directed evolution from an ancestral fungal laccase, catalyzes the one-electron oxidation of diverse aromatic substrates using O_2_ instead of toxic H_2_O_2_, and exhibits activity selective to the surface plasma membrane of both living and fixed cells. We show that LaccID can be used with mass spectrometry-based proteomics to map the changing surface composition of T cells that engage with tumor cells via antigen-specific T cell receptors. In addition, we use LaccID as a genetically-encodable tag for EM visualization of cell surface features in mammalian cell culture and in the fly brain. Our study paves the way for future cell-based applications of LaccID.

## Introduction

Proximity labeling (PL) has emerged as a powerful technique for mapping spatial proteomes, interactomes, and transcriptomes in living cells and animals^1^. Current PL enzymes fall largely into two broad classes: the peroxidases and the biotin ligases (**Fig. 1a**). Peroxidases such as APEX2^2^ and horseradish peroxidase (HRP)^3^ generate phenoxyl radicals from phenol substrates and H_2_O_2_, which subsequently react with surface-exposed electron-rich sidechains like tyrosines. These enzymes are fast and versatile, able to use a wide range of substrates, but the requirement for H_2_O_2_ renders the labeling conditions toxic, especially in tissue. Labeling by biotin ligases such as TurboID^4^ and BioID^5^ is milder; these enzymes use endogenous ATP to generate biotin-AMP esters that subsequently react with lysine sidechains. However, due to the strict requirement of these enzymes for biotin, it is difficult to avoid background from both endogenous biotin and endogenous biotinylated proteins^6, 7^, which are abundant in vivo^8^. Furthermore, biotin ligases have rarely been used for cell surface mapping due to the absence of ATP in the extracellular environment. While ATP could be added exogenously, tissue penetration and unintentional activation of purine receptors are concerns^9, 10^. For these reasons, new PL enzymes that use different chemistries than existing enzymes are needed to advance the field.

**Figure 1.**
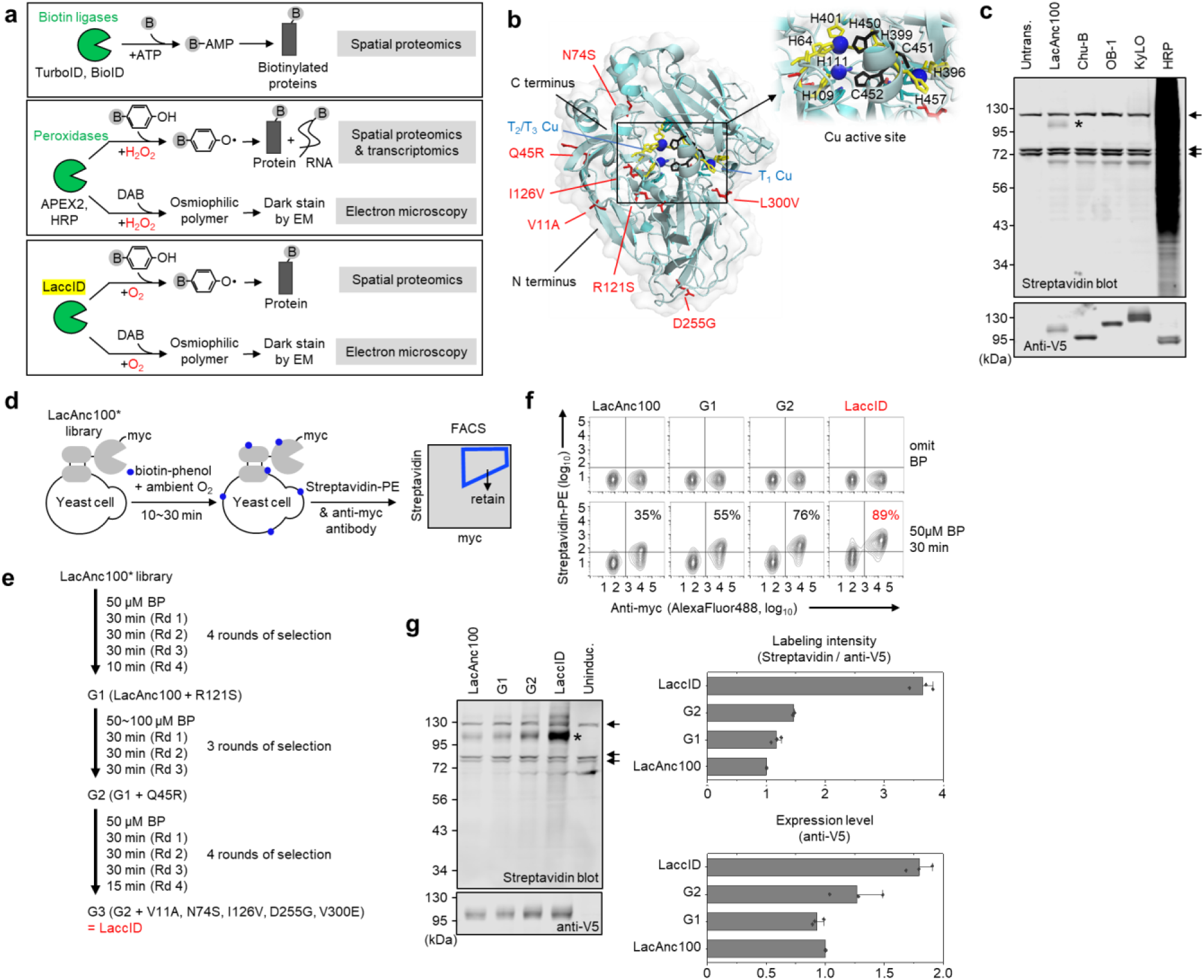
Directed evolution of LaccID. (**a**) LaccID is versatile, like APEX2, because it can be used for proximity labeling as well as electron microscopy (EM). LaccID is also minimally toxic, like TurboID, because it uses O_2_ instead of toxic H_2_O_2_. Grey B, biotin. DAB, diaminobenzidine. (**b**) AlphaFold2-predicted structure of LaccID. Four copper atoms (blue) are distributed in two copper centers (one Cu atom in type I and three Cu atoms in type II/III centers). Cu-coordinating sidechains (Histidines 64, 109, 111, 396, 399, 401, 457) are shown in yellow, residues involved in electron transfer (His_451_, Cys_452_, His_453_) in black, and seven mutations introduced by directed evolution in red. (**c**) Initial screening of 4 laccase candidates: LacAnc100^24^, ChU-B^25^, OB-1^26^, and KyLO^67^. Enzymes were fused to the transmembrane (TM) domain of human CD4 for expression on the surface of HEK293T cells. Labeling was performed for 1 h with 500 μM biotin-phenol (BP), then cell lysates were analyzed by streptavidin blotting. For comparison, HRP labeling was 1 min with 500 μM BP μM and 1 mM H_2_O_2_. *Self-labeling of LacAnc100. Arrows point to endogenous biotinylated proteins. Untrans., untransfected. (**d**) Yeast surface display selection scheme. PE, phycoerythrin. (**e**) Selection conditions used over three generations of directed evolution. The winning clone from each generation was named G1, G2, and G3, respectively. G3 is LaccID. (**f**) FACS density plots summarizing the progress of directed evolution. Yeast cells displaying LacAnc100 or G1-G3 (LaccID) were compared by labelling with 50 μM BP for 30 min, then staining with streptavidin-PE and anti-myc antibody. This experiment was performed three times with similar results. Percentages calculated as fraction of myc+ cells with streptavidin-PE staining above background. (**g**) Comparison of LacAnc100 and evolved clones on the surface of HEK293T cells. Expression of cell surface-targeted constructs was induced with doxycycline overnight, then labeled for 1 h with 500 μM BP in EBSS. Asterisk indicates self-labeling of LaccID. Arrows point to endogenous biotinylated proteins. Uninduc., uninduced. This experiment was performed three times with similar results. Right: quantification of streptavidin blot data (n = 3 biological replicates each; error bars, s.d.).

Given the versatility of phenoxyl radicals for covalent tagging of proteins^11^, RNA^12^, DNA^13^, and for generating contrast for electron microscopy (EM)^14^, we searched for APEX2-like oxidizing enzymes that do not require toxic H_2_O_2_ for labeling. Polyphenol oxidase (PPO)^15^ and laccase^16^ are both copper enzymes that oxidize phenol using O_2_ instead of H_2_O_2_. In particular, laccase drew our attention because: (1) it catalyzes one-electron oxidation to generate short-lived phenoxyl radicals^16^. PPO, on the other hand, carries out two-electron oxidation to generate quinones^15^ which have much longer half-lives and therefore larger labeling radii. (2) Laccase has broad substrate scope, which includes diaminobenzidine (DAB)^17^, a substrate that is used to generate contrast for EM^14^. (3) Laccase’s K_m_ for O_2_, 20-50 μM^18^, is in the range of physiological O_2_ concentrations in tissue (29-64 μM in the human brain for example^19^) suggesting potential *in vivo* applicability. (4) Laccase has been used for cell surface fluorescence imaging, albeit with antibody targeting rather than genetic-encoding^20^. And finally, (5) fungal laccases^21^, which have broader substrate scope and higher redox potential than their bacterial counterparts, have been engineered for numerous industrial applications, including paper-pulp biobleaching, textile dye decolorization, bioremediation, food processing, and biofuel cells^22^.

However, there are significant challenges to the use of laccases for PL technology, most notably because no laccase has previously been expressed and shown to be active in mammalian systems. The enzyme is also complex, with four copper atoms, two or more disulfides, glycosylation, optimal activity at acidic pH (pH 4-5), and an electron transfer pathway that is strongly interrupted by modest concentrations of hydroxyl anions or halides (e.g., NaCl, KCl) in cell culture media^23^. Here we benchmarked several engineered laccase templates^24–27^, identified an ancestral fungal variant with slight activity in HEK293T cells, and performed extensive directed evolution on the yeast surface to improve its activity (**Fig. 1b**). The resulting clone, LaccID, was deployed for both electron microscopy and cell surface proteome mapping in mammalian cells. Our work establishes a new enzyme class for PL with promising properties and paves the way for future improvements.

## Results

### Engineering LaccID

Laccases are widely expressed in fungi, plants, insects, and bacteria^16^, but there is no mammalian counterpart. The natural function of laccases is polymerization/depolymerization of aromatic substrates involved in lignin degradation/biosynthesis, stress defense, pigment synthesis, and sporulation^28^. Laccases possess four copper atoms distributed into two copper centers. Fungal laccases also contain at least two disulfide bonds and are heavily glycosylated^21, 29^. The most extensively engineered laccases for industrial applications are from white-rot fungi, which have been improved for activity in ionic liquids, high and low pH, high temperature, and physiological fluids^28^.

Given the lack of prior reports demonstrating laccase activity in mammalian systems, we first selected a panel of fungal laccase variants to test in HEK293T cells. We cloned them as fusions to a secretion signal, an epitope tag (V5), and transmembrane anchor (from CD4^30^) for cell surface expression. Our panel consisted of: (i) LacAnc100^24^, an ancestrally resurrected sequence from Basidiomycete PM1 fungi, which may offer improved heterologous expression and activity due to the generalist nature of ancient biocatalysis^31^; (ii) Chu-B^25^, a mutant of PM1 laccase that was engineered for applications in human blood, withstanding neutral/alkaline pH and high halide concentrations; (iii) OB-1^26^, another mutant of PM1 laccase engineered for improved expression in budding yeast, suggesting potential compatibility with mammalian expression systems; and (iv) KyLO^27^, a laccase mutant from ascomycetous *Myceliophthora thermophila* engineered for enhanced activity at pH 8.0.

All four laccases were well-expressed in HEK293T cells, as determined by anti-V5 western blotting (**Fig. 1c**). However, only a single laccase, LacAnc100, showed labeling activity when incubated with biotin-phenol (BP) probe for 1 hour. LacAnc100 exhibited a small degree of self-labeling (asterisk in **Fig. 1c**), which was less than the intensity of endogenous biotinylated protein bands and far lower than the promiscuous labeling catalyzed by horseradish peroxidase (HRP) with BP and H_2_O_2_ in just 1 minute. Yet this small activity provides a basis to carry out directed evolution of LacAnc100 to improve its properties.

We implemented directed evolution using yeast surface display (**Fig. 1d**) due to our past success with this format for engineering APEX2^2^ and TurboID^4^. In addition, fungal laccases have been functionally expressed in *S. cerevisiae* at high titers^32–35^, indicating that yeast provide the necessary machinery for both copper insertion and glycosylation^36^. We fused LacAnc100 to the yeast mating protein Aga2p for surface display and used error-prone PCR to create a library of variants with an average of 1-7 amino acid changes per gene.

BP labeling of the yeast library was followed by streptavidin-fluorophore staining and fluorescence-activated cell sorting (FACS) to enrich populations with high activity. Several strategies were employed over 11 rounds of selection. First, we progressively increased selection stringency by reducing labeling time (**Supplementary Figs. 1a, 2a**, and **3a**). Second, at the end of each generation, all unique sequences were tested in HEK293T cells and beneficial mutations were manually combined (**Supplementary Figs. 1-3**) before diversification of the winning clone for further evolution. Third, in some rounds, half of the cell population was subjected to labeling in the presence of the radical quenchers sodium ascorbate and Trolox to reduce radical half-life and intercellular “trans” labeling; the populations were recombined after FACS enrichment (**Supplementary Fig. 2**).

In total, we performed 11 rounds of selection over three generations of evolution (**Fig. 1e**). FACS analysis of labeled yeast showed a gradual increase in biotinylation efficiency over the course of evolution, with LaccID being 4-fold more active than the original template LacAnc100 (**Fig. 1f**). This trend was reproduced when the clones were analyzed on the surface of HEK293T cells (**Fig. 1g**), with LaccID exhibiting both highest expression and the highest activity-to-expression ratio of all clones.

### Optimization of LaccID labeling conditions and characterization in mammalian cells

Although LaccID has higher biotinylation activity than the starting template LacAnc100, the activity is still much lower than that of HRP in a side-by-side comparison (**Supplementary Fig. 4a**). We hypothesized that the mammalian cell environment, including the culture media, may impact LaccID activity. Thus, we tested different media compositions, comparing complete DMEM to RPMI and EBSS media (**Fig. 2a**). We found that labeling with BP was strongest in EBSS, perhaps because it lacks thiols that could inhibit laccase activity^37^. In a separate test, we found that additives such as 10% FBS and 0.5 mM cysteine can impair LaccID activity in EBSS (**Supplementary Fig. 4b**).

**Figure 2.**
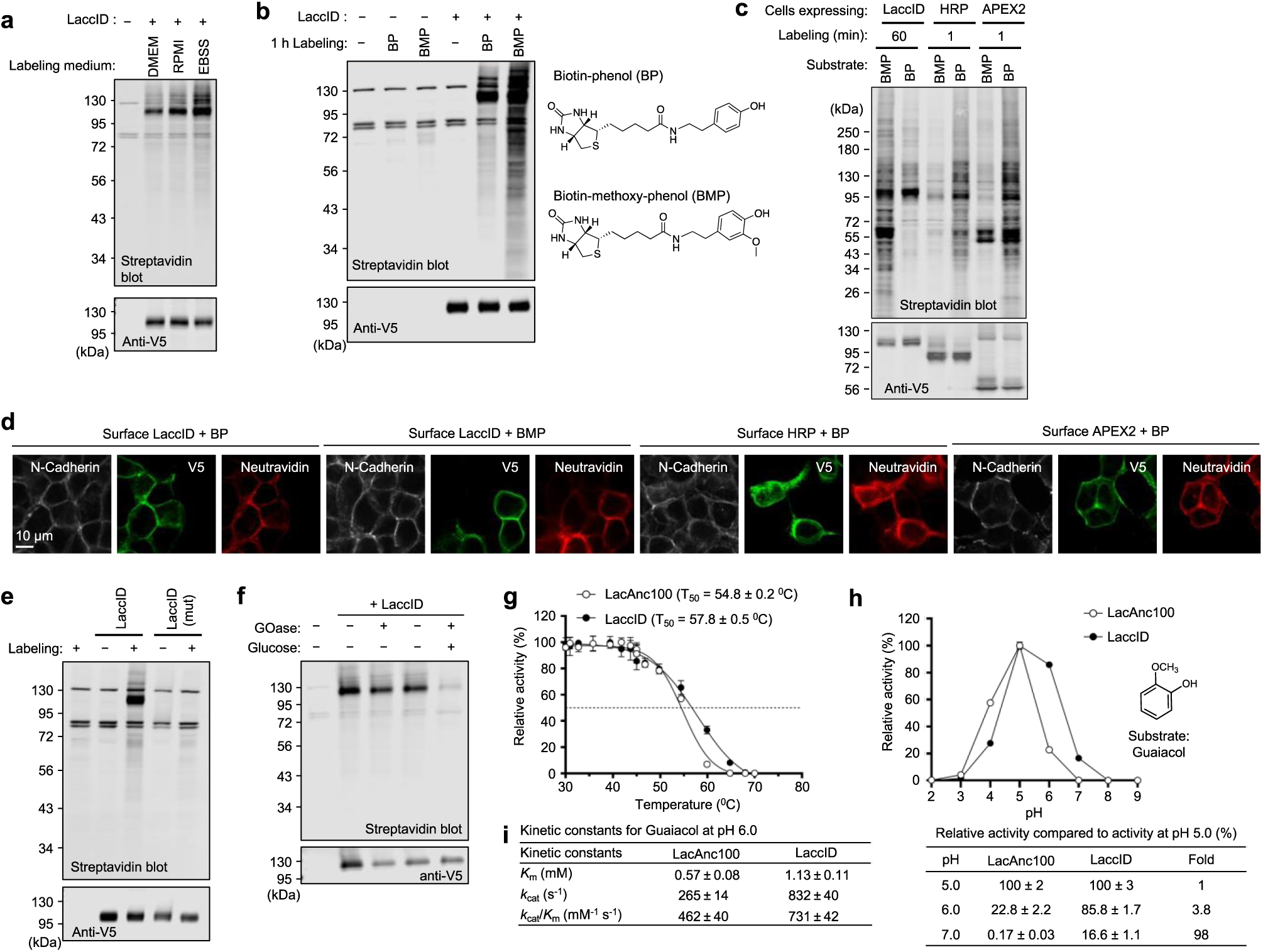
Characterization of LaccID. (**a**) LaccID labeling in different cell culture media. HEK293T cells expressing surface LaccID were labeled for 2 h with 500 μM BP. (**b**) LaccID labeling with BP versus BMP (biotin-methoxy-phenol). HEK293T cells expressing surface LaccID were labeled for 1 h with 500 μM probe in EBSS. This experiment was performed three times with similar results. (**c**) LaccID comparison to HRP and APEX2 on the surface of HEK293T cells. For LaccID, labeling was performed with 500 μM BP in EBSS or 500 μM BMP in RPMI. For HRP and APEX2, labeling was performed with 500 μM BP and 1 mM H_2_O_2_ in DMEM. LaccID molecular weight is 53 kD without glycosylation. (**d**) Confocal imaging of cells labeled as in **c**. N-cadherin stain is shown as a plasma membrane marker. Neutravidin detects biotinylated proteins. Anti-V5 detects enzyme expression. (**e**) Mutation of the HCH motif (His_450_, Cys_451_, His_452_) abolishes LaccID activity. HEK293T cells expressing LaccID or mutant LaccID were labelled for 1 h with 500 μM BP in EBSS. This experiment was performed three times with similar results. (**f**) LaccID requires O_2_. HEK293T cells expressing surface LaccID were labeled with 500 μM BP in PBS for 1 h. Glucose oxidase (GOase) and glucose were used to deplete oxygen in the culture media. (**g**) Thermal stability of LacAnc100 and LaccID. Purified enzymes were incubated in citrate-phosphate pH 6.0 buffer at temperatures ranging from 30 to 70 °C, then assayed for activity at pH 4.0 with ABTS substrate. Three technical replicates each; error bars, s.d. (**h**) pH-activity profile for LacAnc100 and LaccID. Purified enzymes were assayed in Britton and Robinson buffer at the indicated pH with the substrate guaiacol. Conversion to product was measured by Abs_470_. Additional data with other substrates (ABTS, 2,6-dimethoxyphenol, *p*-coumaric acid, and sinapic acid) in **Supplementary Figs. 5c-d**. (**i**) Kinetic constants for LacAnc100 and LaccID at pH 6.0 with guaiacol. Additional data with other substrates in **Supplementary Fig. 5e**.

We next hypothesized that the substrate structure may influence biotinylation by LaccID. Many natural and synthetic substrates used by laccases have an ortho-methoxy substituent on the phenol, an electron-donating moiety that facilitates oxidation^38^. When tested on LaccID-expressing HEK293T cells, biotin-methoxyphenol (BMP) showed higher labeling than BP (**Fig. 2b**), and comparable labeling in both RPMI and EBSS media (**Supplementary Fig. 4c**).

Substrate titration and time-course experiments showed that 1-2 hours of labeling with 250-500 μM BP or BMP substrate provides optimal signal to noise ratio (**Supplementary Figs. 4d-e**). Interestingly, while LaccID shows stronger labeling with BMP than BP, HRP and APEX2 exhibit the opposite preference (**Fig. 2c**). Confocal immunofluorescence imaging confirmed the surface localization and activity of LaccID (**Fig. 2d**). When using BMP, in contrast to BP, cells adjacent to LaccID-expressing cells were also biotinylated to some extent (trans-labeling) (**Fig. 2d**), which was also observed on suspended K562 cells analyzed by FACS (**Supplementary Fig. 4f**). The BMP radical is expected to have a longer half-life than BP radical due to stabilization by the electron-donating methoxy group, thus resulting in a longer labeling radius. Consequently, for PL applications with LaccID, BMP is recommended to maximize sensitivity, and BP to maximize spatial specificity.

To examine the mechanism of LaccID labeling, we generated a mutant in which the His-Cys-His (HCH) motif, essential for electron transport^39^, was changed to three alanines. As expected, this mutation abolished LaccID activity (**Fig. 2e**). To confirm that LaccID labeling depends on O_2_, we depleted O_2_ from the labeling mixture with glucose oxidase, which consumes O_2_ upon oxidation of its substrate glucose^40^. As expected, LaccID labeling decreased when both glucose oxidase and glucose were present **(Fig. 2f** and **Supplementary Fig. 4g**).

### Kinetic characterization of LaccID

To further characterize LaccID’s catalytic activity, especially in relation to the starting template LacAnc100, we purified recombinant enzyme using a protease cleavable His_8_-tag from yeast *S. cerevisiae* (**Supplementary Fig. 5a**). We first performed a thermal stability assay by increasing temperature while measuring oxidation of 2,2’-Azino-bis(3-ethylbenzothiazoline-6-sulfonic acid) (ABTS), a commonly-used laccase substrate^41^. LaccID was more stable than LacAnc100, with a T_50_ (temperature at which the enzyme loses 50% of activity after 10 minutes of incubation) of 57.8±0.5 ℃ compared to 54.8±0.2 ℃ for LacAnc100 (**Fig. 2g**). This aligns with the higher expression of LaccID compared to LacAnc100 observed in HEK293T cells (**Fig. 1g**). There was no significant difference in pH-dependent stability between the two enzymes (**Supplementary Fig. 5b**).

To measure catalytic turnover, we utilized a broader panel of substrates: ABTS, guaiacol, 2,6-dimethoxy phenol (DMP), *p*-coumaric acid, and sinapic acid. Initial rates of product generation were measured by absorbance, showing that LaccID was 1.1 to 20.8-fold faster than LacAnc100 with the same substrates at pH 7.4 (**Supplementary Fig. 5c**). We then measured initial rates across a range of pH values. We observed that while LacAnc100 activity drops at neutral pH (pH 7.0) to only 0.17% of its peak value at pH 5.0, LaccID retains 16.6% of its peak (pH 5.0) activity at pH 7.0 (**Fig. 2h** and **Supplementary Fig. 5d**). We also performed quantitative kinetic measurements at pH 6.0 (pH 7.0 was not possible for technical reasons explained in Methods; **Fig. 2i** and **Supplementary Fig. 5e**) and observed improvements in both k_cat_ and k_cat_/K_m_ for LaccID compared to LacAnc100.

Finally, we measured sensitivity of both laccases to halides, specifically chloride. We did not detect a difference in chloride IC_50_, the concentration at which activity is reduced by 50% (**Supplementary Fig. 5f**). Collectively, our in vitro measurements suggest that the improvements in LaccID result from a combination of higher thermal stability and increased turnover at physiological pH compared to LacAnc100.

To better understand the structural basis for this activity improvement, we used AlphaFold3^42^ to model LaccID and LacAnc100. **Supplementary Fig. 6** suggests that two mutations - N74S and R121S - give rise to structural changes that re-orient the copper-coordinating sidechains His451 and His453.

### LaccID activity is specific to the mammalian cell surface

In mammalian cells, LacAnc100 is active at the cell surface while fusions targeted to the cytosol, mitochondria, or endoplasmic reticulum lumen are inactive (**Supplementary Fig. 7a**). In nature, laccases are processed through the secretory pathway which completes the glycosylation, copper insertion, and disulfide formation necessary to generate active enzyme; these post-translational modifications may not be possible in other compartments of the cell.

To similarly test LaccID, we created an ER-targeted construct by appending a KDEL ER retention sequence, and prepared analogous constructs for APEX2 and HRP for comparison. Immunofluorescence staining showed correct ER localization for all constructs, with high overlap with calreticulin, an endogenous ER marker (**Fig. 3a**). We performed a side-by-side comparison of all ER-targeted constructs with all cell surface targeted constructs (**Fig. 3a-b**). Interestingly, surface-targeted LaccID gave much cleaner localization to the cell surface than surface-targeted APEX2 or HRP, which exhibited significant pools of trapped intracellular protein (**Fig. 3a**). The enhanced surface targeting of LaccID was further confirmed by antibody staining, with and without membrane permeabilization, followed by FACS analysis (**Supplementary Fig. 7b**). We then performed labeling with BP probe. While APEX2 and HRP showed BP labeling on both the cell surface and ER lumen, LaccID was exclusively active on the cell surface, with no labeling detected by ER-targeted LaccID (**Figs. 3a-b**).

**Figure 3.**
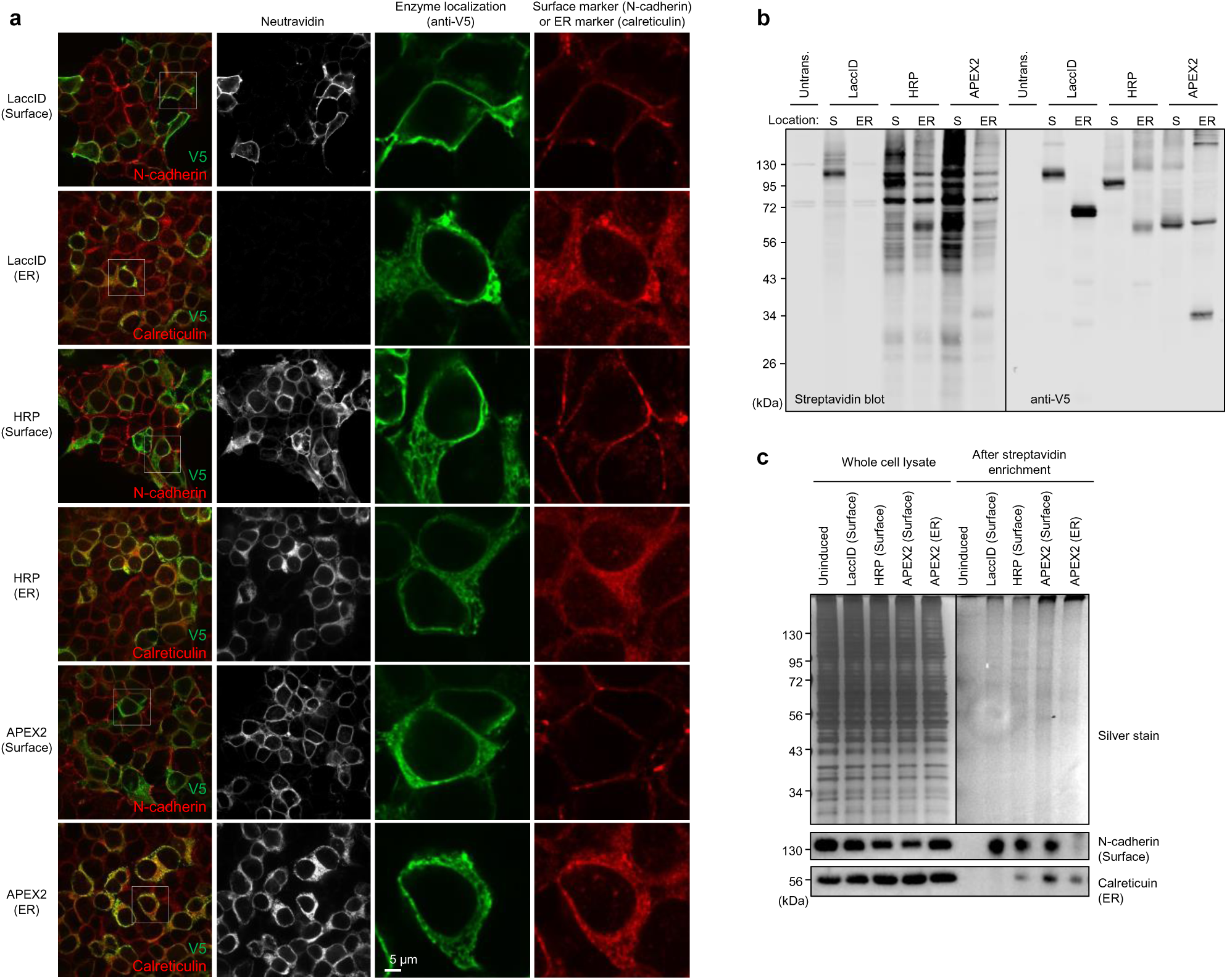
LaccID is selectively active on the cell surface. (**a**) Confocal imaging of enzymes targeted to the cell surface or ER lumen of HEK293T cells. LaccID samples were labeled with BP for 1 h in EBSS. HRP and APEX samples were labeled for 1 min with BP and H_2_O_2_ in DMEM. The cells were then fixed and stained with anti-V5 antibody to detect expression, neutravidin-AF647 to detect biotinylated proteins, and organelle markers as indicated. This experiment was performed twice with similar results. (**b**) Promiscuous biotinylation activity of enzymes at the surface (S) or in the ER lumen of HEK293T cells. Cells were labeled with BP as in **a**, then blotted with streptavidin and anti-V5 antibody. APEX2 molecular weight is 30.9 kD and the higher band may be a crosslinked dimer. (**c**) Western blot detection of proteins enriched by LaccID, HRP, or APEX, targeted to the cell surface or ER lumen. Lysate from HEK293T cells labeled as in **a** were enriched with streptavidin beads, and eluates were blotted for a surface marker protein (N-cadherin) or ER lumen marker protein (calreticulin).

To further examine the cell surface specificity of LaccID, we performed promiscuous labeling with BP, enriched the biotinylated proteins with streptavidin beads, and blotted the enriched material with antibodies against the ER marker calreticulin and the cell surface marker N-cadherin. **Fig 3c** shows that surface-targeted LaccID enriches N-cadherin but not calreticulin, while surface-targeted APEX2 and HRP enrich both markers due to (1) significant pools of APEX2/HRP in the ER lumen due to their imperfect targeting to the cell surface, and (2) high activity of APEX2/HRP in the ER compartment. The serendipitous discovery of LaccID’s cell surface selectivity provides an opportunity for facile cell surface proteome mapping, explored below.

LaccID’s inactivity in the ER may result from limited intracellular copper, which is tightly regulated and facilitated by copper chaperones predominantly in the Golgi^43^. This hypothesis is supported by our observation that ER-targeted LaccID was unable to catalyze diaminobenzidine (DAB) polymerization after cell fixation, but incubation of the fixed cells with CuSO_4_ restored activity (**Supplementary Fig. 7c**). We also generated a construct in which LaccID was targeted directly to the Golgi compartment. Golgi-targeted LaccID failed to show activity with BP in living cells (**Supplementary Fig. 7d**), but it did polymerize DAB in fixed cells (**Supplementary Fig. 7e**). Perhaps the discrepancy is due to lower oxygen availability inside living cells versus after cell fixation. We also examined the importance of glycosylation by introducing Asn-to-Ala mutations at six different potential N-glycosylation sites in surface-targeted LacAnc100, none of which appeared to significantly reduce its activity (**Supplementary Fig. 7f**). Together, our findings indicate that LaccID is well-targeted to the cell surface, and selectively active there, in contrast to APEX2 and HRP which exhibit activity in ER and Golgi compartments. The cell surface-selective activity may be due to copper insertion pathways but not glycosylation.

### LaccID as a genetically encoded tag for electron microscopy

Genetically-encoded labels analogous to GFP for fluorescence microscopy are highly valuable for marking organelles, proteins, and other cellular features in electron microscopy (EM). Because laccases are known to oxidize diaminobenzidine (DAB)^17^, a molecule used to generate contrast for EM^14^, we tested the use of LaccID as an electron microscopy tag (**Fig. 4a**).

**Figure 4.**
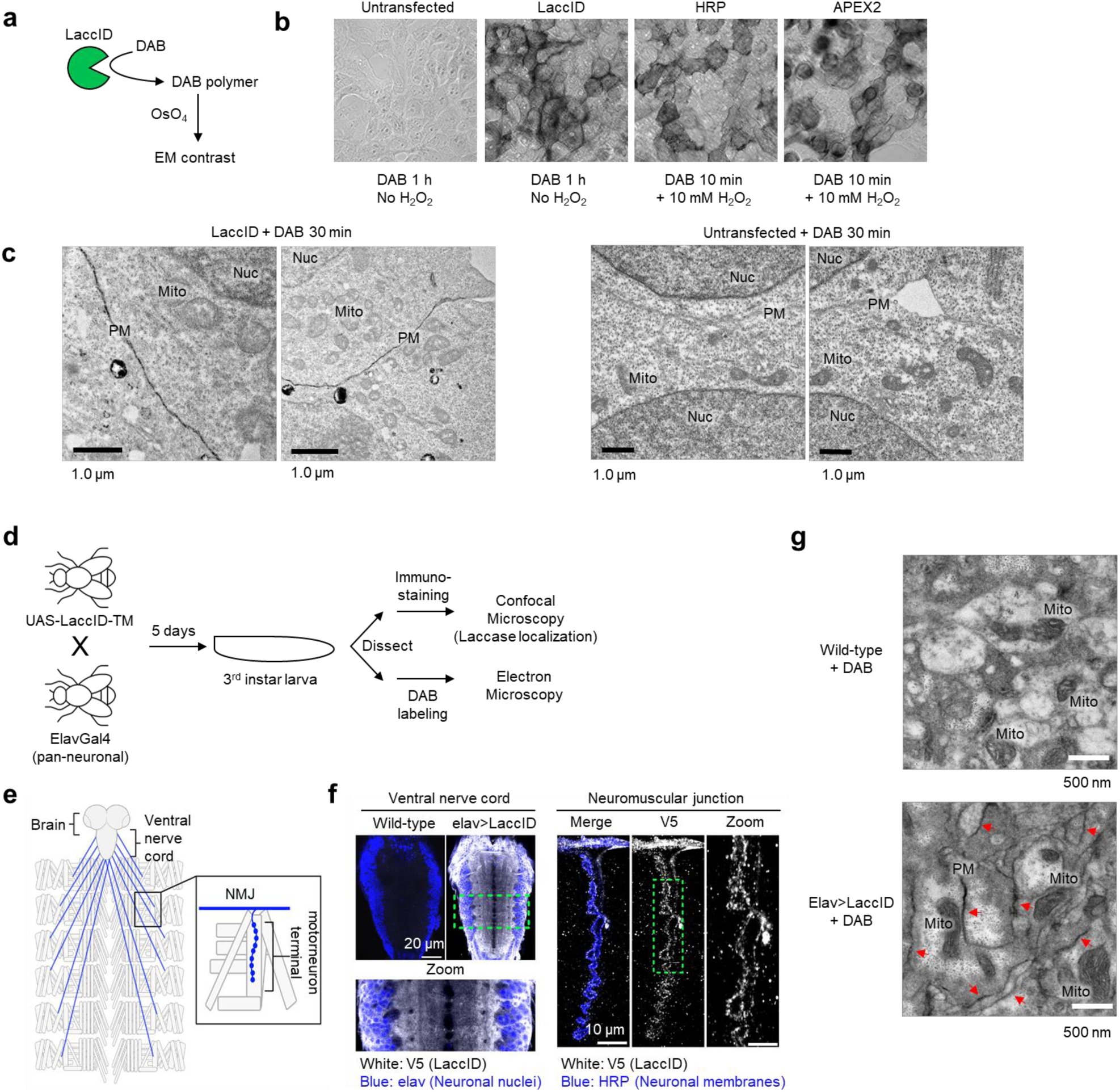
LaccID is a genetically encoded electron microscopy tag. (**a**) LaccID generates EM contrast by catalyzing the oxidative polymerization of DAB, which recruits electron-dense osmium that appears dark on EM micrographs. (**b**) LaccID-catalyzed DAB staining in fixed cells. HEK293T cells expressing surface LaccID, HRP, or APEX2 were fixed using 2% (vol/vol) glutaraldehyde, stained with DAB for indicated times, and imaged by brightfield microscopy. (**c**) EM imaging of HEK293T cells expressing LaccID targeted to the cell surface via fusion to the transmembrane domain of CD4. Cells were fixed using 2% glutaraldehyde, incubated with DAB for 30 min, then post-fixed with 1% osmium tetroxide for one hour. Images are representatives of >3 fields of view. PM, plasma membrane. Nuc, nucleus. Mito, mitochondria. Right: Negative control with LaccID expression omitted. (**d**) Generation of neuron-specific LaccID transgenic flies. TM, transmembrane domain of CD2. (**e**) Schematic of the larval *Drosophila* central nervous system and the innervation of body wall muscles by motor neuron axons. LaccID is expressed in neurons of the brain and ventral nerve cord (VNC). Inset shows motorneuron axons leaving the VNC to innervate muscles at neuromuscular junctions (NMJs), which are visualized by confocal microscopy in **f**. (**f**) Representative confocal images of anti-V5 immunostained LaccID (epitope tagged with V5) in the VNC or NMJ. Neuronal nuclei and membranes were labeled with anti-elav and anti-HRP^68^, respectively. Boxed regions are magnified to show LaccID expression in neuronal cell bodies and neuropil in the VNC (left) and at a motor neuron terminal (right). (**g**) Representative EM images of DAB-stained fly ventral nerve cord (VNC). Fixed VNC tissues from fly larvae were stained with Ce-DAB2 for 75 min, followed by post-fixation with 1% osmium tetroxide for 1 h. Red arrows indicate plasma membrane labeling in LaccID-expressing samples (Elav>LaccID). Additional fields of view and cerium detection by electron energy loss spectroscopy (EELS) and energy-filtered TEM are in **Supplementary Figs. 8c, d, e**.

We prepared HEK293T cells expressing cell surface LaccID, fixed, and overlaid with DAB. LaccID catalyzed the oxidative polymerization and local deposition of DAB, which could be visualized by brightfield microscopy (**Fig. 4b**). Interestingly, the difference in activity compared to cell surface HRP and APEX2 in this context was considerably smaller than the activity difference observed for biotinylation on living cells. Possible reasons could be that (i) DAB is preferred by LaccID over other aromatic substrates, and (ii) DAB labeling was performed in sodium cacodylate buffer which does not contain laccase inhibitors like halides and thiols.

Generating EM contrast at the cell surface delineates cell-cell borders and allows efficient cell segmentation and synapse reconstruction^44^. For such applications, achieving efficient surface labeling while minimizing intracellular labeling is crucial. We hypothesized that LaccID might provide more selective cell surface labeling than peroxidases, based on the observations above (**Fig. 3**). TEM imaging of OsO_4_-postfixed HEK293T cells shows that surface LaccID provides excellent contrast at the plasma membrane, with minimal staining of intracellular membranes on the ER, mitochondria, and nucleus (**Fig. 4c**). By contrast, APEX2, also targeted to the cell surface, provided much weaker staining of the plasma membrane relative to intracellular organelles (**Supplementary Fig. 8a-b**).

Next, we tested whether we could extend LaccID-based EM imaging to the fruit fly *Drosophila* melanogaster. We generated transgenic flies expressing LaccID in neurons by crossing *UAS-LaccID* flies with *elav-GAL4* (pan-neuronal GAL4) flies (**Fig. 4d**). Confocal immunofluorescence imaging of the fly larval ventral nerve cord (VNC) confirmed neuron-specific expression of LaccID, and its cell surface localization at axon termini at neuromuscular junctions (**Figs. 4e-f**). TEM imaging of fixed VNCs shows dark contrast at the plasma membrane when both LaccID and Cerium-DAB2 are present (**Fig. 4g** and **Supplementary Fig. 8c**). In **Supplementary Fig. 8d**, cerium elemental maps show that the dark-stained regions in EM are indeed positive for cerium atoms, which are coordinated by the DAB-chelate conjugate^45^. These findings indicate that LaccID can serve as a peroxide-free, surface-specific, genetically encodable EM marker suitable for use in fixed cells and fly tissues.

### LaccID for mapping cell surface proteomes

Cell surface proteomes (“surfaceomes”) drive development, intercellular communication, synapse formation, immune responses, and a vast array of other biological processes in multicellular organisms. Methods to map cell surface proteomes in an unbiased, comprehensive, and cell type selective manner are increasingly needed to probe such biology. Peroxidase-mediated PL of surface proteomes has been used in fly^46^ and mouse^8^ brains to discover novel proteins that mediate nervous system development. However, these studies have been limited by the toxicity of H_2_O_2_ and the poor tissue permeability of BxxP, a membrane-impermeant variant of BP required to prevent labeling of intracellular ER and Golgi proteins by surface-targeted HRP^3^. Because LaccID does not require H_2_O_2_, and its activity is inherently selective for the cell surface (obviating the need for special probes such as BxxP), we wondered if LaccID could be used for nontoxic mapping of cell type specific surfaceomes in complex biological settings.

We established a co-culture assay to examine the interactions between human primary CD8+ T cells and tumor cells expressing a specific tumor antigen, NY-ESO-1^47^. The T cells were engineered to express cell surface LaccID and either a T cell receptor (TCR) or chimeric antigen receptor (CAR) against NY-ESO-1 (**Supplementary Fig. 9a**). The T cells were then cocultured with NY-ESO-1+ A375 tumor cells in a 3:1 ratio for 24 hours before labeling with BMP for 2 hours (**Fig. 5a** and **Supplementary Fig. 9b**).

**Figure 5.**
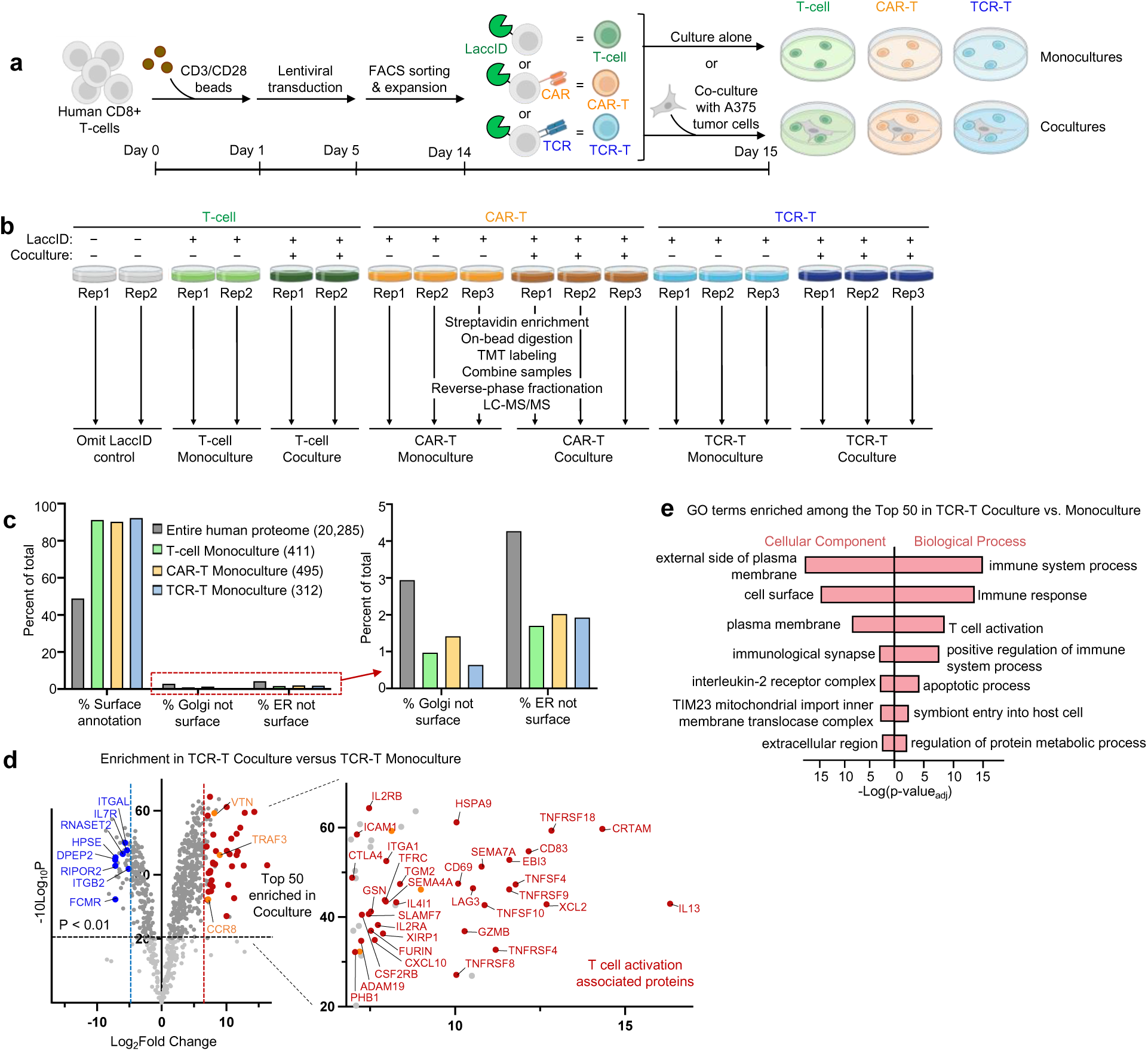
LaccID mapping of the T cell surface proteome in the presence of tumor cells. (**a**) Cocultures were established between LaccID-expressing T cells and A375 melanoma tumor cells. The tumor cells were positive for the NY-ESO-1 tumor antigen, while T cells expressed a CAR (orange) or TCR (blue) directed towards the same antigen, or neither (green). (**b**) Design of 18-plex TMT proteomic experiment. All samples were treated with BMP for 2 hours prior to quenching, lysis, and streptavidin enrichment of LaccID-labeled proteins. (**c**) LaccID enriches cell surface proteins in all three monoculture samples. Golgi and ER-resident proteins (lacking cell surface annotation) are de-enriched. (**d**) Volcano plot comparing LaccID-labeled TCR-T cocultures to monocultures. Proteins with prior literature connections to T cell activation are colored red, those with indirect connections are colored orange, and previously-annotated downregulated proteins are colored blue. Details in **Supplementary Table 3**. (**e**) Gene ontology terms enriched in top 50 TCR-T coculture vs. monoculture dataset.

18 samples were quantitatively compared via tandem mass tag (TMT)-based mass spectrometry proteomics (**Fig. 5b**). We observed a high correlation between biological replicates (**Supplementary Fig. 9c**), and good segregation of samples by coculture status and receptor expression via principal component analysis (**Supplementary Fig. 9d**). To obtain cell surface proteome lists for control T-cell, CAR-T cell, and TCR-T cell monocultures (grown in the absence of tumor cells), we applied a log_2_-fold change cut-off with a 5% false discovery rate (FDR), defined by the retention of cytosolic and nuclear proteins, which should not be labeled by LaccID (**Supplementary Fig. 9e**). This resulting protein lists (312 to 495 in size; **Supplementary Table 2** and **Supplementary Fig. 9f-g**) were highly enriched in cell surface proteins, with 90-92% of entries having prior cell surface annotation (**Fig. 5c** and **Supplementary Fig. 9f**). Notably, few ER-resident and Golgi-resident proteins were enriched in any of the proteomes (**Fig. 5c**), consistent with our observation that LaccID lacks activity in these compartments.

Coculture surfaceomes were determined in a similar manner, but due to the higher presence of intracellular proteins (cytoskeletal proteins in the T-cell and CAR-T cell cocultures, and mitochondrial proteins in the TCR-T cell cocultures), we applied a less stringent cutoff (10% instead of 5% false discovery rate, **Supplementary Fig. 10a**). The intracellular proteins could reflect transfer from tumor cells to immune cells via extracellular vesicles, mitochondria-plasma membrane interactions^48^, or tunneling nanotubes^49^, but they could also be non-specific binding artifacts due to our use of the monoculture (rather than coculture) no-LaccID negative control for data filtering (**Fig. 5b**). The final coculture protein lists (450-595 in size) are 76-81% specific for cell surface proteins (**Supplementary Table 2** and **Supplementary Fig. 10b**).

Comparison of monoculture and coculture datasets showed little evidence of T cell activation for the T-cell (no TCR and no CAR) samples, as expected (**Supplementary Figs. 10c, e**). The CAR-T cells showed activation (e.g., upregulation of CD40LG, IL2RA, and CSF2RB markers) even in the absence of tumor cells, contrary to TCR-T cells (**Supplementary Figs. 9g-i**), consistent with previous reports of antigen-independent signaling by CARs^50^. Upon coculture, CAR-T cell surfaceomes did not change much (**Supplementary Figs. 10d, f**), because the cells were already activated, and this CAR may exhibit low antigen sensitivity^51^. The greatest change was observed for TCR-T cells, which strongly upregulated many known markers of T cell activation upon coculture with tumor cells. For example, the activation markers CD69, CD83^52^, LAG3^53^, IL2RA/B, TFRC, TNFRSF4/9/18^54^, and CTLA4^55^ were highly enriched (**Fig. 5d-e**). Among the top 50 hits – proteins most enriched by LaccID in TCR-T cell coculture versus monoculture – were 33 proteins with direct literature links to CD8+ T-cell activation; another 3 proteins with indirect links; 2 membrane proteases (ADAM19 and FURIN); and 8 mitochondrial proteins (**Fig. 5d** and **Supplementary Table 3**). Additionally, we observed downregulation of several markers known to be decreased in activated T-cells, including DPEP2, FCMR, RIPOR2, HPSE, and RNASET2 (**Fig. 5d** and **Supplementary Table 3**).

A few unexpected proteins were present in our TCR-T cell top-50 list, raising the possibility of novel biological hypotheses. For example, prohibitin1 (PHB1) is a ubiquitous protein expressed in multiple subcellular regions, but primarily in the inner mitochondrial membrane^56^. Perhaps our detection of PHB1 in TCR-T cell coculture reflects tumor-induced relocalization of PHB1 from mitochondria^57, 58^ or other intracellular stores to the cell surface. This alteration could be driven by phosphorylation^59^ and could play a role in integrating immune signaling with metabolism^60^. Another protein we identified is gelsolin (GSN), a Ca^2+^-dependent actin binder that severs actin filaments. A recent report showed that GSN is secreted via exosomes from ovarian cancers and can induce T cell inhibition and apoptosis^61^. A375 melanoma cells used in our experiments express GSN^62^. Perhaps our detection of GSN in all co-cultures relative to monocultures reflects GSN secretion from A375 tumor cells to facilitate immune evasion^61, 63^.

Our findings indicate that LaccID can be a useful tool for mapping endogenous cell surface proteomes in complex samples, in a non-toxic (H_2_O_2_-free) and cell-type specific manner.

## Discussion

We have engineered LaccID as a new oxidizing enzyme for proximity labeling and electron microscopy applications. LaccID offers unique benefits over established PL enzymes such as APEX2 and TurboID by utilizing gentle, non-toxic (H_2_O_2_-free) labeling conditions, and accepting a broad range of substrate structures. In addition, LaccID can be used for selective proteome mapping at the cell surface without requiring special, membrane-impermeant substrates. This enabled us to use LaccID to map the surfaceome of T cells cocultured with tumor cells, and highlight dozens of proteins that are upregulated via T cell receptor-antigen interactions (**Fig. 5**). We also used LaccID for EM imaging in the fly brain and HEK293T cell cultures, delineating plasma membranes more clearly than peroxidase-based labels such as APEX2 (**Fig. 4** and **Supplementary Fig. 8**). These examples showcase the promise of this new class of PL enzyme.

LaccID was created from an ancestral fungal laccase via three generations of yeast display directed evolution, which improved enzyme stability, catalytic turnover, and pH-activity profile to better suit physiological conditions (**Figs. 1, 2**). However, the current activity of LaccID is still much lower than that of APEX2 or TurboID, requiring 1-2 hours of labeling to give detectable signal, in contrast to APEX2 or TurboID which need only 1-10 minutes. Future generations of LaccID may benefit from directed evolution in mammalian cells, which have different glycosylation and copper insertion machinery than yeast. In addition, selection in the presence of serum, halides, and thiols could help to enrich variants that are tolerant to those additives. Our current LaccID did not exhibit activity in living fly brain tissue, but an improved version with in vivo activity would open the door to many PL applications.

Apart from LaccID, several antibody-based, non-genetically-encoded methods have recently been reported for mapping cell surface proteomes in culture. For example, transition metal^64^ and organic dye^65^ photocatalysts have been conjugated to antibodies against cell surface markers to profile the microenvironment of the EGF receptor^64^, the surfaceome of erythrocytes^66^, and the proteome of Jurkat cell-HEK293T cell immune synapses^65^. The main drawbacks of such methods compared to LaccID are the difficulty of achieving cell type-specific profiling if suitable surface markers (that can be targeted by antibodies) are unavailable (by contrast, LaccID expression can be controlled by cell-type specific promoters and viruses), and, for future applications, the challenge of delivering large antibody reagents as well as light into animals and tissue.

A major goal of connectomics is to use EM to accurately trace neurons by distinguishing cell surface membranes from intracellular membranes in thin tissue sections. LaccID provides a powerful tool with inherent selectivity for the surface membrane based on its biochemical mechanism.

Beyond PL and EM, LaccID may have utility in other areas. Engineered fungal laccases have been used for drug, food, and paper production, as well as for removal of toxic substances such as herbicides and industrial effluents^22^. The improved thermal stability and activity of LaccID at neutral pH could serve useful in these settings. Our work paves the way for novel biological and biotechnological applications of this versatile enzyme class.

## Methods

### Cloning

All constructs were generated using standard cloning techniques, as previously described^6^. Briefly, gene fragments were obtained from Integrated DNA Technologies or Twist Biosciences, or amplified using Q5 polymerase (NEB Cat# M0491S). Vectors were digested using enzymatic restriction digest and ligated to either gel-purified PCR products or ordered gene fragments using Gibson assembly. Ligated plasmid products were transformed into competent XL1-Blue *E. coli*. Detailed information on the constructs is provided in the **Supplementary Information**.

### Mammalian cell culture, transfection and generation of stable cell lines

Mammalian cell cultures were performed following previously published protocols^6^. Briefly, HEK293T cells (ATCC, CRL-3216) were cultured in complete DMEM (Dulbecco’s modified Eagle’s medium (DMEM, Gibco Cat# 11965-092) supplemented with 10% (v/v) fetal bovine serum (FBS, VWR Cat# 97068-085) and 1% penicillin-streptomycin (VWR Cat# 16777-164)) at 37 °C under 5% CO_2_. For transient expression, cells in 12-well plates were typically transfected at roughly 70% confluency using 2.5 μL of polyethyleneimine (PEI, 1 mg/mL in water, pH 7.3) or Lipofectamine 2000 (Life Technologies) and a total 1,000 ng of plasmid in 100 μL of serum-free media. Stable cell lines were generated with lentivirus infection. Lentivirus was generated by transfecting HEK293T cells plated to roughly 60% confluency in a six-well dish with 250 ng of pMD2.G, 750 ng of psPAX2 and 1,000 ng of lentiviral vector containing the gene of interest with 10 μL of PEI in serum-free media. Lentivirus-containing supernatant was collected after 48 h and filtered through a 0.45 μm filter. HEK293T cells at roughly 60% confluency were infected with crude lentivirus followed by selection with 10 μg/mL puromycin for 1 week. To induce expression of proteins under doxycycline-inducible promoter (pTRE3G), cells were incubated with media supplemented with 100–200 ng/mL doxycycline overnight.

### Labeling in mammalian cells with LaccID

HEK293T cells expressing LaccID or its variants were prepared through doxycycline induction of stable cells or PEI transfection of LaccID-encoding plasmids. To perform labeling, cell culture media was replaced with media containing 500 μM of probe and incubated at 37 °C. Best LaccID labeling conditions are: 500 μM BP for 1-2 h in EBSS, or 500 μM BMP for 1-2 h in RPMI. Labeling with HRP and APEX2 was performed as previously described^3^. After labeling, cells were quenched with 10 mM sodium ascorbate and 5 mM Trolox in DPBS, washed three times more with DPBS, and analyzed by Western blot or immunofluorescence as described below.

### LaccID labeling with various additives

For oxygen depletion experiments, labeling was performed in the presence of 4% (w/v) glucose and 0.25 mg/mL glucose oxidase (Sigma-Aldrich, Cat# G2133). For BP labeling with different additives, each component was added with following concentrations: 5, 10, 15% FBS, 0.5, 1, 1.5% penicillin-streptomycin, 50, 100, 150 mM CaCl_2_, and 0.03, 0.06, 0.09 g/L cysteine, glycine, or serine.

### Western blotting

Western blotting was performed as previously described^6^. Briefly, after labeling and washing, cells were lysed with RIPA lysis buffer supplemented with 1× protease inhibitor cocktail (Thermo Scientific Cat# 78429). Lysates were cleared via centrifugation at 20,000*g* at 4 °C for 10 min. Cleared lysates were mixed with protein loading buffer and boiled at 95 °C for 5 min. Samples were then loaded on 10% SDS–PAGE gels, transferred onto nitrocellulose membrane, and stained with Ponceau S (Sigma) to visualize total protein loading. Blots were blocked in either 5% (w/v) nonfat milk (Lab Scientific, M-0841) or 1% (w/v) bovine serum albumin (BSA) in 1× tris-buffered saline with Tween (TBST) (Teknova Cat# T1688) for 30–60 min at room temperature, incubated with primary antibodies in TBST for 1 h at room temperature, washed three times with TBST for 5 min each, incubated with fluorophore-conjugated secondary antibodies for 30 min at room temperature, washed three times with TBST and imaged on an Odyssey CLx imager (LI-COR) or ChemiDoc XRS+ imager with Clarity Western ECL Blotting Substrates (Bio-Rad). List of antibodies and dilution rates are provided in the **Supplementary information**, “**Antibodies**” section. Quantification of labeling was performed with Image Studio (LI-COR).

### Immunofluorescence imaging

For immunofluorescence imaging, cells were grown on 12 mm-diameter glass coverslips (VWR Cat# 72230-01) in 24-well plates. Cells expressing LaccID were prepared as described above. After labeling, cells were gently quenched, washed three times with DPBS, and were fixed and permeabilized with ice-cold methanol for 10 min. Cells were washed again three times with DPBS and blocked for 1 h with 1% bovine serum albumin (BSA, Fischer Scientific Cat# BP1600-1) in TBST at 4 °C. Cells were then incubated with primary antibodies in TBST overnight at 4 °C In After washing three times with TBST, cells were incubated with fluorophore-conjugated secondary antibodies in TBST for 1 h at room temperature. Cells were washed three times with TBST and imaged.

Imaging was performed with a Zeiss Axio Observer.Z1 microscope with a Yokogawa spinning disk confocal head, Cascade IIL:512 camera, a Quad-band notch dichroic mirror (405/488/568/647 nm), and 405 nm, 491 nm, 561 nm, and 640 nm lasers (all 50 mW). Images were captured through a 63× oil-immersion objective for the following fluorophores: DAPI (405 laser excitation, 445/40 emission), AlexaFluor 488 (491 laser excitation, 528/38 emission), PE and AlexaFluor 568 (561 laser excitation, 617/73 emission), and AlexaFluor 647 (647 laser excitation, 700/75 emission). Image acquisition times ranged from 10 to 500 ms per channel, and images were captured as the average of two or three such exposures in rapid succession. Image acquisition and processing was carried out with the SlideBook 5.0 software (Intelligent Imaging Innovations, 3i).

### Yeast cell culture

Yeast cell cultures were performed as previously described^6^. Briefly, *Saccharomyces cerevisiae* strain EBY100 was propagated at 30 °C in SDCAA media supplemented with 20 mg l^−1^ tryptophan. Competent yeast cells were transformed with yeast-display plasmid pCTCON2 following the Frozen EZ Yeast Transformation II (Zymo Research Cat# T2001) according to the manufacturer’s protocol. Successful transformants were selected in SDCAA plates and propagated in SDCAA media. To induce protein expression, yeasts were inoculated from saturated cultures in 10% SD/GCAA (SDCAA with 90% of dextrose replaced with galactose) supplemented with 100 μM CuSO_4_ overnight.

### Analysis of LaccID activity on yeast surface

Yeast cells expressing surface-displayed LaccID mutants as Aga2p fusions in a pCTCON2^6^ vector were cultured overnight in SD/GCAA media to induce expression. Next, 6 × 10^6^ cells (1 OD_600_ = 1 × 10^7^ cells) were transferred into PBS with 0.1% BSA (PBS-B), containing 50-100 μM of BP. Cultures were then incubated at 30 °C or 37 °C while rotating. After BP labeling, cells were washed with 10 mM sodium ascorbate and 5 mM Trolox in PBS-B. Yeast cells were washed three times more in cold PBS-B by pelleting at 3,000x*g* for 2 min at 4 °C and resuspending in 1 ml cold PBS-B. Yeast cells were stained in 50 μL of PBS-B containing anti-Myc antibody (1:400) for 1 h at 4 °C for detecting enzyme expression, then the cells were washed three more times and stained in 50 μL of PBS-B with goat anti-chicken-AF488 and SA-PE for 40 min at 4 °C. Samples were washed for the final three times with 1 ml PBS-B before analysis or sorting. To analyze and sort, single yeast cells were gated on a forward-scatter area (FSC-A) by side-scatter area (SSC-A) plot around the clustered population (P1) on a ZE5 Cell Analyzer (Bio-Rad). P1 was then gated on a side-scatter width (SSC-W) by side-scatter height (SSC-H) plot around the clustered population (P2). P2 populated cells were then plotted to detect AF488 and phycoerythrin signals. FlowJo v10 (BD Biosciences) was used to analyze FACS data.

### Yeast directed evolution

Yeast directed evolution was performed following previously published protocols^6^. In each generation, LacAnc100 mutant library was generated using error-prone PCR with 100 ng of template plasmid. In each generation, three libraries were generated at different levels of mutagenesis with the following primers:

Fwd: 5′-ggaggaggctctggtggaggcggtagcggaggcggagggtcggctagccatatg-3′

Rev: 5′-gatctcgagctattacaagtcctcttcagaaataagcttttgttcggatcc-3′

PCR products were gel purified then reamplified for 30 more cycles under normal conditions and gel purified again. The pCTCON2 backbone was digested with NdeI and BamHI and gel purified as well. Both 4,000 ng of PCR product and 1,000 ng of cut vector were mixed and dried in a DNA speed vacuum (DNA110 Savant). The dried DNA was then resuspended in 10 μL of water and electroporated into electrocompetent EBY100 yeasts. After electroporation, yeasts were rescued in 2 ml of yeast extract peptone dextrose media and recovered at 30 ℃ without shaking for 1 h. Then, 1.98 mL of the culture was propagated in 100 mL of SDCAA medium while 20 μL was used to determine library size. Yeasts were diluted 100×, 1,000×, 10,000×, 100,000× and 20 μL of each dilution was plated onto SDCAA plates at 30 ℃ for 3 days. The resulting library size was determined by the number of colonies on the 100×, 100×, 1,000× and 100,000× plates, corresponding to 10^4^, 10^5^, 10^6^ or 10^7^ transformants in the library, respectively. Libraries with different mutation levels were combined in equal volume after testing the enzyme activity, to form the yeast-display library.

### Directed evolution of LaccID: generation 1

For the first round of evolution, three libraries were generated using LacAnc100 as the starting template. The three libraries were generated using error prone PCR as described above, using the following conditions to produce varying levels of mutagenesis:

Library 1: 2 μM 8-oxo-dGTP, 0.5 μM dPTP, 20 PCR cycles

Library 2: 2 μM 8-oxo-dGTP, 1 μM dPTP, 20 PCR cycles

Library 3: 2 μM 8-oxo-dGTP, 2 μM dPTP, 20 PCR cycles

The library sizes (approximated by number of transformants as described above) were 6×10^7^ for each library. FACS analysis of the three libraries showed robust expression and wide range of activities for all three libraries. All libraries were combined (named E1) and used as the initial population for the first round of selections. This combined library was induced as described above.

Population E1 was passaged twice, and from this culture approximately 1.1 x 10^9^ cells were prepared for sorting (assuming 1 OD_600_ ≈ 1 × 10^7^ cells) as described above with 50 μM BP labeling for 30 min. 4.8×10^8^ cells were sorted by FACS. A triangular gate that collected cells positive for both anti-myc and streptavidin was drawn, and approximately 4×10^6^cells were collected (∼0.8%) to give population E1R1. Subsequent rounds were performed in a similar manner, and the details of each round are as following:

| Library | Cells prepared | Labeling | Labeling time (min) | Cells sorted by FACS | Cells collected (%) |
| --- | --- | --- | --- | --- | --- |
| E1R1 | 1.5 × 10<sup>8</sup> | 50 μM BP | 30 | 4.0×10<sup>7</sup> | 1.7×10<sup>4</sup> (<0.1%) |
| E1R2 | 1.5 × 10<sup>8</sup> | 50 μM BP | 30 | 2.9×10<sup>7</sup> | 9.8×10<sup>3</sup> (<0.1%) |
| E1R3 | 7.5 × 10<sup>7</sup> | 50 μM BP | 10 | 1.4×10<sup>7</sup> | 9.2×10<sup>3</sup> (<0.1%) |
| E1R4 | 7.5 × 10<sup>7</sup> | 50 μM BP | 10 | 1.4×10<sup>7</sup> | 5.0×10<sup>3</sup> (<0.1%) |

After five rounds of enrichment, 24 individual clones were sequenced from E1R4 and E1R5, and unique sequences were tested in HEK293T cell surface. Clones with higher activity (quantified by streptavidin signal/anti-V5 signal) were chosen. Combining mutations in higher-activity clones didn’t result in increased activity. Clone with an R121S mutation showed the best activity and was named G1. See Supplementary Figure 1 for more details.

### Directed evolution of LaccID: generation 2

For the second round of evolution, three libraries were generated using G1 (LacAnc100 + R121S) as the starting template. The three libraries were generated using error prone PCR as described above, using the following conditions to produce varying levels of mutagenesis:

Library 1: 2 μM 8-oxo-dGTP, 2 μM dPTP, 20 PCR cycles

Library 2: 4 μM 8-oxo-dGTP, 1 μM dPTP, 20 PCR cycles

Library 3: 8 μM 8-oxo-dGTP, 0.5 μM dPTP, 20 PCR cycles

The library sizes (approximated by number of transformants as described above) were 6×10^7^ for each library. FACS analysis of the three libraries showed robust expression and wide range of activities for all three libraries. All libraries were combined (named E2) and used as the initial population for the first round of selections. This combined library was induced as described above.

Starting from E2R1, the directed evolution was performed in two parallel branches (for detail, see Supplementary Figure 2). In the first pathway, LaccID expression level was quantified by staining with anti-chicken-AF647. In the second pathway, staining was performed with anti-chicken-AF488, to prevent bleed-over of PE signal to AF647 channel. In addition, in round 4 and 5 sorting of the second pathway, two reactions were performed in parallel: labeling in the absence/presence of low-concentration quenchers to prevent transcellular labeling. Libraries from parallel labeling were combined after checking the activity of each population. As previously described, after each round a triangular gate that collected cells positive for both anti-myc and streptavidin was drawn for sorting. Details of each round are as following:

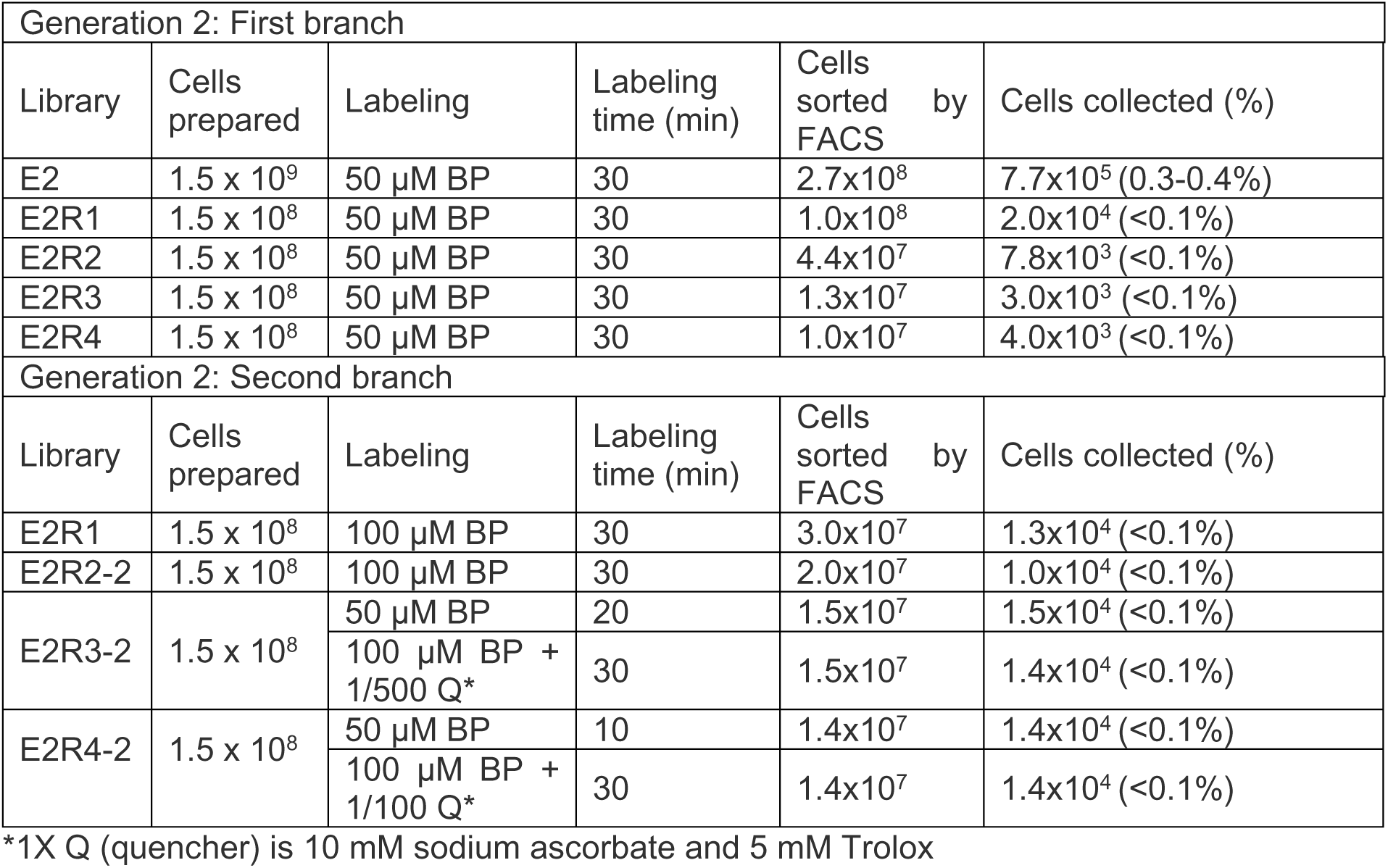

20 individual clones were sequenced from E2R4, E2R5, E2R3-2, E2R4-2, and E2R5-2. Unique sequences were tested in HEK293T cell surface. Clones with higher activity (quantified by streptavidin signal/anti-V5 signal) were chosen. Combining mutations in higher-activity clones didn’t result in increased activity. Clone with a Q45R mutation showed the best activity and was named G2. See Supplementary Figure 2 for more details.

### Directed evolution of LaccID: generation 3

For the third round of evolution, three libraries were generated using G2 (LacAnc100 + Q45R, R121S) as the starting template. The three libraries were generated using error prone PCR as described above, using the following conditions to produce varying levels of mutagenesis:

Library 1: 30 μM 8-oxo-dGTP, 1 μM dPTP, 20 PCR cycles

Library 2: 20 μM 8-oxo-dGTP, 2 μM dPTP, 20 PCR cycles

Library 3: 10 μM 8-oxo-dGTP, 3 μM dPTP, 20 PCR cycles

The library sizes (approximated by number of transformants as described above) were 6×10^7^ for each library. FACS analysis of the three libraries showed robust expression and wide range of activities for all three libraries. All libraries were combined (named E3) and used as the initial population for the first round of selections. This combined library was induced as described above. Details of each round are as following:

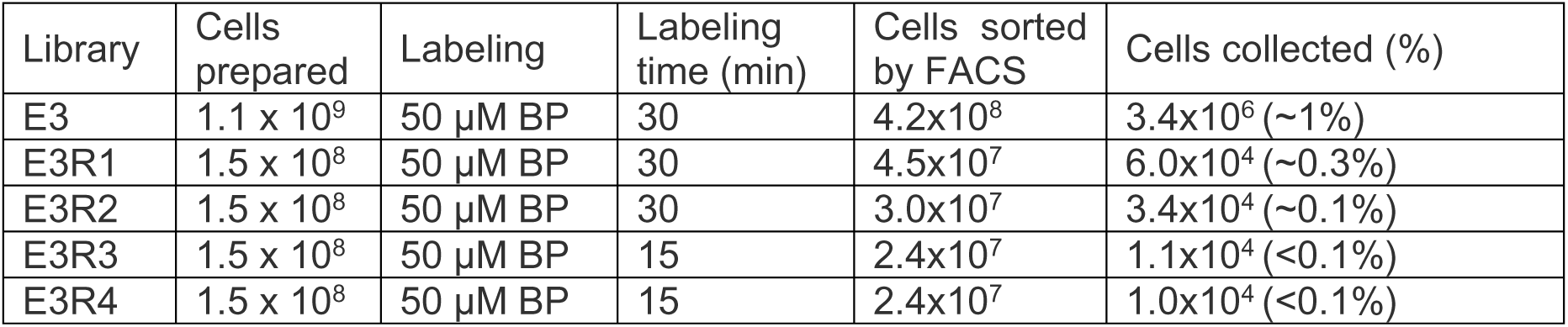

20 individual clones were sequenced from E3R3, E3R4, and E3R5. Unique sequences were tested in HEK293T cell surface. Clones with higher activity (quantified by streptavidin signal/anti-V5 signal) were chosen. Among 4 clones that showed high activity, combining mutations in 3 clones resulted in an increased activity. The mutant with G2 mutations + mutations from all 3 clones was named G3, which is subsequently named LaccID. See Supplementary Figure 3 for more details.

### K562 cell culture and coculture trans-labeling experiments

K562 cells were cultured in RPMI1640 media supplemented with 10% (v/v) FBS and 1% penicillin-streptomycin at 37 °C under 5% CO_2_. Cells were routinely passaged in fresh cell culture media to keep the cell density under 1.0 x 10^6^ cells/mL. Cell lines stably expressing cytosolic GFP and surface LaccID were achieved by lentivirus infection, followed by FACS sorting.

For BMP labeling, cells were harvested from culture media and resuspended in RPMI containing 500 μM BMP with a final density of 9.0 x 10^5^ cells/mL. After 2 h labeling at 37 ℃, cells were washed and stained following the same protocol used for yeast labeling. For coculture experiments, 1:1 mixture of wild-type K562 cells and GFP+ cells expressing surface LaccID were labeled and analyzed in the same manner.

### Labeling in mammalian cells with LaccID

HEK293T cells expressing LaccID or its variants were prepared through doxycycline induction of stable cells or PEI transfection of LaccID-encoding plasmids. To perform labeling, cell culture media was replaced with media containing 500 μM of probe and incubated at 37 °C. Best LaccID labeling conditions are: 500 μM BP for 1-2 h in EBSS, or 500 μM BMP for 1-2 h in RPMI. Labeling with HRP and APEX2 was performed as previously described^3^. After labeling, cells were quenched with 10 mM sodium ascorbate and 5 mM Trolox in DPBS, washed three times more with DPBS, and analyzed by Western blot or immunofluorescence as described below.

### Cloning of His-tagged laccase variants for large-scale production

DNA sequences encoding LacAnc100 and LaccID were *in vivo* cloned into *S. cerevisiae* strain BJ5465. To promote *in vivo* recombination in yeast, the genes were PCR-amplified to introduce specific overhangs of 40-55 bp to the linearized vector using iProof High-Fidelity Polymerase (BioRad, #1725301) following manufactureŕs specifications. The PCR products were gel extracted, purified and cleaned (Macherey-Nagel, #740609), and 200 ng of each laccase mutant gene were transformed along with 100 ng of linearized vector into *S. cerevisiae* using yeast transformation kit (Merck, #YEAST1). The linearized vector was a modified version of pJRoC30 vector (Kindly donated by F.H. Arnold, CALTECH) containing a C-terminal HRV-3C protease cleavage site (3C-tag: LEVLFQGP) followed by a GSG linker and 8xHis-tag. Transformed cells were plated on SDCAA drop-out plates and incubated for 3 days at 30 °C. Subsequently, the selected clones were subjected to screening for laccase activity in 96-well microtiter plates before confirmation of plasmids sequence.

Individual clones containing LacAnc100 and LaccID were inoculated into 50 μL of SC minimal medium supplemented with 6.7 g/L yeast nitrogen base (BD, #291940), 1.92 g/L yeast synthetic drop-out medium supplement without uracil (Merck, #Y1501), 20 g/L D-raffinose (Merck, #R0250), and 25 μg/mL chloramphenicol (Merck, #C0378)) in a sterile 96-well plate (Enzyscreen, #CR1496c). In the plate, two columns were inoculated with LacAnc100 and LaccID, and two additional columns were used for a negative control plasmid (encoding for a non-related protein). The plates were sealed with AeraSeal films (Merck, # A9224) to allow gas exchange in cultures and incubated at 30 °C, 225 rpm and 80% relative humidity in a humidity shaker (Minitron, INFORS). After 48 h, 160 μL of expression medium (10 g/L yeast extract (Gibco, #212750), 20 g/L peptone (Gibco, #211677), 20 g/L D-galactose (Panreac, #A1131), 67 mM potassium phosphate monobasic buffer pH 6.0 (Thermo Scientific, #424200250), 25 g/L ethanol (Scharlau, #ET00062500), 2 mM copper sulfate (Merck, #C7631), and 25 μg/mL chloramphenicol (Merck, #C0378)) were added to each well. The plate was further incubated for 24 h. Then, the plate was centrifuged for 15 min at 4,000x*g* and 4 °C (Eppendorf 5810R) and replicated with the help of a liquid handler robotic station (Freedom Evo, TECAN) by transferring 20 μL of supernatant into a new replica plate. Next, 180 μL of reaction buffer containing 100 mM citrate-phosphate buffer pH 4.0 and 1 mM ABTS (Panreac, #A1088) as a substrate were added using a Multidrop robot (Multidrop Combi, Thermo Scientific). The plate was stirred briefly, and laccase activity was determined following the absorption of ABTS oxidation at 418 nm (εABTS^•+^ = 36,000 M^-1^ cm^-1^) in kinetic mode using a plate reader SpectraMax ABS Plus (Molecular Devices).

An aliquot from the wells containing the most active LacAnc100 and LaccID clones was inoculated in 3 mL of YPD (10 g/L yeast extract (Gibco, #212750), 20 g/L peptone (Gibco, #211677), 20 g/L D-glucose (Thermo Scientific, #410955000), and 25 μg/mL chloramphenicol (Merck, #C0378)) and incubated at 30 °C and 225 rpm for 24 h. Next, recovery of laccase-containing plasmids from these cultures was performed using Zymoprep Yeast Plasmid Miniprep Kit (Zymo research, #D2001) regarding the manufactureŕs specifications. Zymoprep product containing shuttle vectors was transformed into *E. coli* XL1-Blue competent cells (Stratagene) and plated on LB/amp plates. Single bacterial colonies were inoculated into 5 mL of LB/amp media and grown overnight at 37 °C with 250 rpm shaking (Minitron, INFORS). The plasmids were then extracted by NucleoSpin Plasmid kit (Macherey-Nagel, #740588) and sent for sequencing (Eurofins genomics). Plasmids encoding for laccase variants were sequenced with primers: RMLN (5’-CCTCTATACTTTAACGTCAAGG), RMLC (5’-GCTTACATTCACGCCCTCCC), Lac_seq 1F (5’-ACAACTTCACTGTCCCGGAT), and Lac_seq 2F (5’-ACCCTACAACGACCCAAACA). Verification of laccase sequences was carried out using Snapgene v6 software.

### Recombinant laccase variants expression and purification

A single colony from *S. cerevisiae* transformed with LacAnc100 or LaccID was used to inoculate 30 mL of SC minimal medium supplemented with raffinose (in a 100 mL flask) and incubated for 48 h at 30 °C and 220 rpm (Orbitron shaker, INFORS). Then, the OD600 of these pre-cultures was measured, and fresh cultures were prepared in 120 mL of SDCAA medium in a 250 mL flask (OD600= 0.3). They were incubated until two growth phases had been completed (10-12 h, OD600 = 1) and thereafter, 50 mL of pre-cultures were induced with 450 mL of expression medium (10 g/L yeast extract (Gibco, #212750), 20 g/L bacto peptone (Gibco, #211677), 20 g/L D-galactose (Panreac, # A1131), 67 mM potassium phosphate monobasic buffer pH 6.0 (Thermo Scientific, #424200250), 25 g/L ethanol (Scharlau, #ET00062500), 2 mM copper sulfate (Merck, #C7631), and 25 μg/mL chloramphenicol (Merck, #C0378)) in a 2.5 liter baffled flask (Quimigen, #931136-B)(OD600= 0.1). The cultures were incubated for 5 days at 30 °C and 130 rpm (Multritron, INFORS) using AeraSeal films (Merck, #A9224) on top of the flask. Next, the cells were pelleted by centrifugation at 4,500x*g* for 10 min at 4 °C (High-Speed Centrifuge CR22N, Eppendorf). The supernatant was collected and doubly filtered (with a glass filter then with a PVDF filter). Clarified cell-free extracts were purified by IMAC (immobilized-metal affinity chromatography). Ni-NTA agarose resin (Qiagen, #30210) was equilibrated in buffer A (Tris·HCl: 20 mM pH 8.0, 150 mM NaCl) and His-tagged laccases were bound to the resin. Non-specific bound proteins were washed out with buffer A supplemented with 25 mM imidazole (Thermo scientific, #A10221) and laccases were eluted with buffer A containing 250 mM imidazole. The eluted proteins were dialyzed against buffer A using an Amicon Ultra-15 device with a molecular cut-off of 10 kDa (Millipore, #UFC901008) and concentrated to 0.5 mL. Next, proteins were treated with 3C-HRV protease (Genescript, #Z03092) overnight at 4°C without shaking according to manufacturer instructions. The following day, Ni-NTA resin equilibrated in buffer A was added over the cleaved laccase samples and incubated at 25 °C and 1,000 rpm in a thermomixer comfort shaker (Eppendorf) for 4 hours. Samples were spun down at 1,000xg for 5 min at 4 °C and the supernatant containing tag-free laccases was collected. Resin containing His-tagged 3C-HRV, non-cleaved laccases and cleaved tags was discarded. Laccase purity and integrity were analyzed by SDS–PAGE using TGX 4-20% Precast gels (Bio-Rad, #4561094) and protein concentrations were estimated using the Bio-Rad protein reagent adapted to 96-well microtiter plates (BioRad, #5000006). Purified laccases were stored at 4 °C for further analysis.

### Kinetic parameters

Steady-state kinetics constants were estimated with purified laccase variants for ABTS (Panreac, #A1088), DMP (2,6-dimetoxyphenol, Merck, #D135550), Guaiacol (Thermo scientific, #120192500), and Sinapic acid (Apollo scientific, #BIS8112) in 100 mM sodium phosphate-citrate buffer pH 5.0 or 6.0. We were unable to measure the kinetic parameters at pH 7.0 due to the impractical amount of recombinant enzyme required for the assay. The concentrations for ABTS ranged from 0.001 to 10 mM, for DMP from 0.02 to 24 mM, for Guaiacol from 0.2 to 60 mM and for Sinapic acid from 0.025 to 0.35 mM. The data was fitted to a non-linear substrate inhibition model (Y = V_max_*X/(Km + X*(1+X/K_i_)). The catalytic efficiency (k_cat_/K_m_) was obtained by plotting turnover rates (s−1) vs. substrate concentrations and fitting it to the equation Y = catEff*X/(1+InvKm*X*(1+X/Ki)), where “catEff” is the catalytic efficiency and “InvKm” is 1/K_m_. Enzymatic reactions were performed in 96-well plate format (Greiner Bio-One, #655101) with 190 µL of reaction mix and 10 µL of enzymes diluted in 10 mM sodium phosphate buffer pH 7.0 at the following concentrations: 1.36 µg/mL (LacAnc100) and 0.81 µg/mL (LaccID) for ABTS, 1.7 µg/mL (LacAnc100) and 0.81 µg/mL (LaccID) for DMP and Guaiacol, and 6.8 µg/mL (LacAnc100) and 8.1 µg/mL (LaccID) for Sinapic acid. For measurements with Sinapic acid 96-well plates were UV-Transparent microplates (Corning, #3635). Reactions were run at room temperature in triplicate and the following molar absorption coefficients were used: ABTS, ε_418nm_ = 36,000 M^−1^ cm^−1^; DMP, ε_469nm_ = 27,500 M^−1^ cm^−1^; Guaiacol, ε_470nm_ = 12,100 M^−1^ cm^−1^; Sinapic acid, ε_312nm_ = 17,600 M^−1^ cm^−1^; and *p*-coumaric acid, ε_300nm_ = 17,200 M^−1^ cm^−1^. Kinetic constants were determined using 65 kDa for molecular weight of laccase estimated by MALDI-TOF^24^. Data was analyzed with GraphPad Prism (v9.5.1) and represented as the mean ± standard deviation.

### Thermostability (T_50_)

Kinetic thermostability of purified LacAnc100 and LaccID were estimated by assessing their T_50_ values using 96-well gradient thermocyclers (T100 thermal cycler, BioRad). Appropriate dilutions of enzymes were prepared such that 20 μL aliquots of enzymes in reaction mix produced a linear response in kinetic mode in the spectrophotometric read. A temperature gradient was set in the thermocycler for LacAnc100 and LaccID ranging from 30 to 70 °C as follows (in °C): 30.0, 30.9, 32.7, 35.8, 39.4, 41.8, 43.7, 45.0, 46.8, 49.8, 54.4, 59.9, 64.8, 68.0, and 70.0. Then, 50 μL of enzymés dilutions were incubated for 10 min at each temperature point into non-skirted 96-well PCR plates (VWR, #732-2387) sealed with thermoresistant adhesive sheets (Thermos scientific, #AB-0558). After this 10 min incubation, samples were immediately chilled out on ice for 10 min and further incubated at room temperature for 5 min. Then, the PCR plates were briefly spun down in a PCR-plate centrifuge (VWR) and 20 μL were transferred into a new flat bottom 96-well plate with the help of a liquid handler (Freedom Evo, TECAN). The laccase activity was measured by adding 180 μL of reaction mixture (100 mM citrate-phosphate buffer pH 4.0 and 1 mM ABTS) by using a Multidrop robot (Multidrop Combi, Thermo Scientific). The plates were briefly stirred, and activity was monitored at 418 nm in kinetic mode using a plate reader SpectraMax ABS Plus (Molecular Devices). The maximum laccase activity detected at 32.7 °C was defined as 100% (initial activity), and other temperature points were normalized to that activity (relative activity). The T_50_ value was defined as the temperature that reduces laccase activity by 50% of its initial activity after 10 min of incubation. Thermostability reactions were measured in three biological replicates. Data was analyzed with GraphPad Prism (v9.5.1) and represented as the mean ± standard deviation (in percentage).

### pH-activity profiles

Appropriate dilutions of purified laccase variants were prepared such that 20 µL aliquots of enzymes in reaction mix gave rise to a linear response in kinetic mode for each substrate assayed. Reaction mixtures were prepared in 100 mM Britton and Robinson buffer at different pH values (2.0, 3.0, 4.0, 5.0, 6.0, 7.0, 8.0, and 9.0) containing ABTS (1 mM), DMP (5 mM), Guaiacol (50 mM), Sinapic acid (0.35 mM), or *p*-coumaric acid (0.1 mM) in each case. One hundred ninety microliters of reaction mixtures were added into flat-bottom 96-well plates (Greiner Bio-One, #655101), except for Sinapic acid and *p*-coumaric acid measurements that were in UV-Transparent microplates (Corning, #3635), with a Multidrop robot (Multidrop Combi, Thermo Scientific). Plates were preloaded with 10 µL of laccase dilutions prepared in 20 mM citrate-phosphate buffer pH 7.0 as follows: 0.28 µg/mL (LacAnc100) and 0.36 µg/mL (LaccID) for ABTS, 1.4 µg/mL (LacAnc100) and 1.8 µg/mL (LaccID) for DMP, 3.4 µg/mL (LacAnc100) and 4.0 µg/mL (LaccID) for Guaiacol, 1.7 µg/mL (LacAnc100) and 8.1 µg/mL (LaccID) for Sinapic acid, and 340 µg/mL (LacAnc100) and 405 µg/mL (LaccID) for *p*-coumaric acid. The plates were briefly stirred and activity was spectrophotometrically determined (SpectraMax ABS Plus, Molecular Devices) at room temperature in kinetic mode at the specific wavelength for each substrate. The relative activities at each pH value (expressed as percentages) were normalized to the maximum activity observed for each variant in the assay. Data was analyzed using GraphPad Prism (v9.5.1) and represented as the mean ± standard deviation (in percentage) of three biological replicates.

### pH-stability profiles

Appropriate dilutions of purified laccase variants were prepared such that 20 µL aliquots of enzymes in reaction mix gave rise to a linear response in kinetic mode. Five microliters of laccase variants (3.4 µg/mL of LacAnc100 and 4.0 µg/mL of LaccID) were incubated in 1 mL final volume containing 100 mM Britton and Robinson buffer at different pH values (2.0, 3.0, 4.0, 5.0, 6.0, 7.0, 8.0, and 9.0) in a 96-deep well plate (Sarstedt, #82.1972) covered with a lid. The plate was stirred and stored in fridge at 4 °C during the experiment. Aliquots of 20 µL were taken from every pH value at different time points (1, 4, 48, 96 and 144 hours) and added over 180 µL of 20 mM Britton and Robinson buffer at pH 7.0, which was preloaded in a bottom flat 96-well microtiter plate (Greiner Bio-One, #655101). The plate was stirred and 20 µL of the mix were transferred to a new 96-well microtiter plate. Then, 180 µL of reaction mix (100 mM citrate-phosphate buffer pH 4.0 and 1 mM ABTS) were added by using a Multidrop robot (Multidrop Combi, Thermo Scientific) and laccase activity was measured at room temperature in a plate reader SpectraMax ABS Plus (Molecular Devices) at 418 nm. Laccase activity at every time point was normalized to the activity value of each variant at time 1 hour at pH 8.0, where the maximum laccase activity was observed. Data was analyzed using GraphPad Prism (v9.5.1) and represented as the mean ± standard deviation (in percentage) of three technical replicates.

### Determination of Initial Turnover numbers (ITN)

The initial turnover rates were determined in phosphate buffer saline at pH 7.4 (PBS, 8 g/L NaCl (Labkem, #SOCH-00T), 1.44 g/L Na_2_HPO_4_ (Supelco, #1.06586), 0.2 g/L KCl (Panreac, #131494), 0.24 g/L KH_2_PO_4_ (Fisher scientific, #BP362)) using ABTS (1 mM), DMP (5 mM), Guaiacol (50 mM), Sinapic acid (0.35 mM), or *p*-coumaric acid (0.1 mM) as substrate. Twenty microliters of LacAnc100 or LaccID were added to bottom flat 96-well plates (Greiner Bio-One, #655101) and 180 µL of reaction mixtures containing PBS and different substrates were added with the help of a Multidrop system (Multidrop Combi, Thermo Scientific). For Sinapic acid and *p*-coumaric acid measurements 96-well plates were UV-Transparent microplates (Corning, #3635). Plates were stirred and laccase activity was assayed at room temperature in a plate reader SpectraMax ABS Plus (Molecular Devices). The absorbance was followed during initial time points, where the reaction was still linear for every substrate. ITN were defined as µmol of product x µmol enzyme^-1^ x min^-1^ using 65 kDa for molecular weight of laccase estimated by MALDI-TOF^24^. Data was analyzed using GraphPad Prism (v 9.5.1) and represented as the mean ± standard deviation of three technical replicates.

### Inhibition the in presence of chloride

The inhibitory effect of chloride on laccase activity was estimated using ABTS at the optimum pH activity for this substrate, thus pH 4.0. Inhibition was determined by the IC_50_ value, which represents the chloride concentration that reduces 50% the initial laccase activity. Laccase activities were measured using 190 µL of reaction mixtures (100 mM citrate-phosphate buffer pH 4.0, 1 mM ABTS) in the presence and the absence of a gradient of NaCl (5-190 mM), and 10 µL of purified LacAnc100 (0.34 µg/mL) or LaccID (0.40 µg/mL) in bottom flat 96-well plates (Greiner Bio-One, #655101). The plates were briefly stirred, and the absorbance monitored in kinetic mode in a plate reader SpectraMax ABS Plus (Molecular Devices). The activity of laccase in the presence of NaCl was normalized to the activity in the absence of NaCl for each laccase variant. Data was analyzed using GraphPad Prism (v9.5.1) and plotted as the mean ± standard deviation (in percentage) of three technical replicates.

### Preparation of DAB-stained HEK293T cells for electron microscopy

LaccID expressing cells were fixed using warm 2% glutaraldehyde in buffer (100 mM sodium cacodylate with 2 mM CaCl_2_, pH 7.4), then quickly moved to ice. After 1h, cells were washed 5 x 2 min on ice with cold buffer and block for 5 min with cold 20 mM glycine in cold buffer. A freshly diluted solution of 0.5 mg/mL DAB tetrahydrochloride in buffer was added to cells for at least 30 min at room temperature. The cells were then washed 5 x 2 min with cold buffer.

HRP or APEX2 expressing cells were fixed and washed as described above. A freshly diluted solution of 0.5 mg/mL DAB tetrahydrochloride with 10 mM H_2_O_2_ in buffer was added to cells for 1-10 min at room temperature.

### DAB staining and preparation of cultured cells for electron microscopy

The fixative was removed, and cells were washed (5 x 1 min) with 0.1 M sodium cacodylate (Ted Pella, 18851) buffer at pH 7.4 containing 2 mM CaCl_2_ on ice and blocked with 20 mM glycine in 0.1 M sodium cacodylate buffer pH 7.4 containing 2 mM CaCl_2_ for 20 min on ice. Cells with APEX2 or HRP were enzymatically reacted with 2.5 mM DAB (3,3’-diaminobenzidine, Sigma-Aldrich, D8001-10G) with 4 mM H_2_O_2_ (from 30% stock) in 0.1 M sodium cacodylate buffer at pH 7.4 for 3-20 min until the desired brown intensity color of the precipitate was visible. Cells with laccase were enzymatically reacted with DAB in 0.1M sodium cacodylate buffer at pH 7.4 for 3-20 min until the desired brown intensity color was achieved). The cells were then washed (5 x 2 min) with 0.1 M sodium cacodylate buffer, pH 7.4 on ice and were post-fixed with 1% osmium tetroxide (EMS, 19150) in 0.1 M sodium cacodylate buffer, pH 7.4 for 30 min on ice. Post-fixative was removed, and cells were washed (5 x 1min) with 0.1 M sodium cacodylate buffer pH 7.4 containing 2 mM CaCl_2_ on ice and (5 x 1 min) with cold ddH_2_O on ice. Cells were dehydrated with ethanol series 20, 50, 70, 90 and 100% on ice for 1 min each, 100% dry ethanol at room temperature for 3 x1 min, infiltrated with 50:50 dry ethanol:Durcupan ACM resin (Sigma-Aldrich, 44610) for 30 min, then 4 changes of Durcupan at one hour each and embedded in a vacuum oven at 60°C for 72 hours.

### Drosophila stocks and generation of UAS-HA-V5-LaccID flies

Flies were maintained on standard cornmeal media with at 12 hr light-dark cycle at 25 °C.

UAS-HA-V5-LaccID flies were generated from gBlocks containing a wingless signal peptide upstream of two epitope tags (HA and V5) followed by Drosophila codon optimized LaccID fused to CD2. These gBlocks were cloned into a 10x pUASt-attB vector, validated by Sanger sequencing, and injected into embryos bearing an attP40 landing site. G0 flies were crossed to a white– balancer, and all white+ progeny were individually balanced.

### Drosophila immunofluorescence and image acquisition

Wandering third instar larvae (L3) were obtained by crossing a 2:1 ration of adult virgin *Elav-Gal4* females to adult UAS-HA-V5-LaccID males. Wandering L3 larvae were dissected in cold PBS and fixed in 4% PFA for 20 min. Following fixation, L3 filets were washed in PTX (PBS and 0.02% Triton X-100) and blocked in PBT (PBS, 0.02% Triton X-100, and 1% NGS) on a nutator and incubated with primary antibodies for either 2 hr at RT or overnight at 4 °C with agitation. The following concentrations of primary antibodies were used: mouse anti-V5 (1:100; R960-25, Thermo Fisher); rat anti-elav (1:100; Developmental Studies Hybridoma Bank). The following species-specific secondary antibodies were used: Alexa Fluor 488 at 1:300. Alexa Flour 594-conjugated anti-HRP (Jackson Immunoresearch Laboratories, Inc.) was used to label motor neuron membranes at 1:300.

Following immunostaining, L3 CNS or body walls were mounted in SlowFade Gold (Invitorgen) and fluorescent images were acquired on a LSM 980 (Carl Zeiss) using a 20X objective (for the CNS) or 40X/63X objective (for the NMJ).

### Preparation of Ce-DAB2-stained fly tissues for electron microscopy

Fly brains and spinal cords were dissected according to previously described methods. Briefly, brains and spinal cords were rapidly dissected in cold 0.15 M sodium cacodylate pH 7.4 containing 4% paraformaldehyde (PFA) and 0.5% glutaraldehyde. After dissection, tissues were fixed for 2-3 h in the same fixative on ice. Tissues were then put into 0.15 M sodium cacodylate pH 7.4 containing 2.5% glutaraldehyde and fixed for additional 20 min at 4 ℃. Unreacted glutaraldehyde was quenched by incubating the tissues in 50 mM glycine in 0.15 M cacodylate buffer for 20 min at 4 ℃. After quenching, tissues were gently washed 5 times with 0.15 M cacodylate buffer.

### Ce-DAB2 staining and preparation of *Drosophila* larval CNS for electron microscopy

The dissected flies’ central brain and ventral nerve cord were rinsed (5 x 3 min) with 0.15 M sodium cacodylate buffer pH 7.4 containing 2 mM CaCl_2_ on ice and blocked with 20 mM glycine, 50 mM potassium cyanide and 50 mM aminotriazole in 0.15 M sodium cacodylate buffer pH 7.4 containing 2 mM CaCl_2_ for 20 minutes on ice. Flies were then washed (2 x 1 min) with 0.15 M sodium cacodylate buffer pH 7.4 containing 2 mM CaCl_2_ on ice and were reacted with 2.5 mM Ce-DAB2 solution in cacodylate buffer^45^ for 75 min to produce the desired brown intensity color from the precipitate. Flies were washed (5 x 3 min) with 0.15 M sodium cacodylate buffer pH 7.4 containing 2 mM CaCl_2_ on ice, post-fixed with 1% osmium tetroxide in 0.15 M sodium cacodylate buffer, pH 7.4 for 1 hour on ice, washed (5 x 3 min) with ddH_2_O on ice, dehydrated in ethanol series 20, 50, 70, 90 and 100% for 5 min each at on ice, 100% dry ethanol at room temperature for 2 x 5 min, 1:1 v/v dry ethanol: dry acetone for 5 min at room temperature and dry acetone for 5 minutes at room temperature.

Flies were infiltrated with the following: 25% Durcupan epoxy resin to 75% dry acetone for 4 hours, 50% Durcupan epoxy resin to 50% dry acetone overnight, 75% Durcupan epoxy resin to 25% dry acetone for 8 hours, 90% Durcupan epoxy resin to 10% dry acetone for overnight and Durcupan ACM (4x 8-12 hours) and were embedded in a vacuum oven at 60°C for a week.

### Transmission Electron Microscopy

100 nm thick sections were cut by an Ultra 45° Diatome diamond knife using a Leica Ultracut UCT ultramicrotome. Both laccase positive and negative flies were cut in the coronal plane from dorsal to ventral. Sections were picked up on a 50-mesh copper grid (Ted Pella, G50) and were carbon coated on both sides by a Cressington 208 carbon coater to prevent charging of the plastic which can cause drift and thermal damage.

TEM images of laccase in plasma membranes in both cultured cells and flies were acquired with a FEI Spirit at 80kV. Fly images were taken posterior and lateral from median line regions adjacent to ventral nerve chord.

### Electron energy loss spectroscopy (EELS) and Energy-filtered transmission electron microscopy (EFTEM)

Both EELS and EFTEM were performed with an in-column Omega filter JEOL JEM-3200EF TEM operating at 300 KV with a LaB6 electron source. The samples were pre-irradiated for about 30 min at a low magnification to stabilize the sample and minimize contamination. Both the electron energy-loss spectrum, zero-loss images and energy filtered elemental maps were acquired using an Ultrascan 4000 CCD detector from Gatan. The EELS acquisition was 60 seconds long using a 4k x 4k format with a 180 eV slit. The cerium elemental map was obtained at the M5,4 core-loss edge, the onset of which occurs at 883 eV. The EFTEM images of the pre- and post-edges were obtained using a slit of 35 eV width and 20 data sets were collected for each pre and post edge window, respectively at 815 eV, 855 eV and 899 eV. For each pre and post window, 20 images were corrected for x-rays, aligned, stacked and sum stack by z projection using Fiji software. Power-law background subtraction was used to create elemental maps of cerium. The elemental map merge and overlay on the TEM image was accomplished as previously reported.

### Primary human T cell isolation and culture

Human T cells were isolated and cultured as previously described^69^. Briefly, primary CD8+ T cells were isolated from blood of anonymous donors by negative selection (STEMCELL Technologies, #17953). T cells were cultured in human T cell medium (HTCM) consisting of X-Vivo 15 (Lonza, #04-418Q), 5% Human AB serum, and 10 mM neutralized N-acetyl L-Cysteine (Sigma Aldrich, #A9165) supplemented with 30 units/mL IL-2 (PeproTech, #200-02) at 37 °C under 5% CO_2_.

### Engineering of primary human T cells and in vitro killing assay

Engineered TCR and CAR receptors recognize the NY-ESO-1_157-165_ peptide presented by HLA-A*02:01. The engineered TCR used for this experiments, 1G4, is an enhanced affinity TCR previously reported^70^. The engineered CAR used the scFv regions previously described^71, 72^ but uses CD8a hinge/TM domains and 4-1BB costimulatory domain. Both engineered receptors have a Myc tag in the C-term to allow detection of receptor surface expression by flow cytometry.

Lentivirus was produced by transfecting HEK239T cells plated in a T75 flask with 6 μg of psPAX2, 3 μg of pMD2.G, and 12μg of lentiviral vector containing the LaccID, TCR, or CAR with 60 μl of polyethyleneimine (PEI Max 1mg/ml in water, pH 7.3). Lentivirus was collected at 24 and 48 h post transfection and filtered through a 0.22 or 0.45 μm filter. 30 mL of lentivirus supernatant was concentrated via centrifugation at 20,000g at 4°C for 2 h, then resuspended in 600 μl StemMACS HSC Expansion media (Miltenyi Biotec). The concentrated lentivirus was used immediately or stored at −80 °C.

Isolated T cells were activated with Human T-Activator CD3/CD28 Dynabeads (Life Technologies #11131D) at a 1:1 cell-to-bead ratio during 24 h before transduction. Activated T cells were transduced with concentrated virus in StemMACS and HTCM supplemented with IL-2 was replenished at 24 h post-transduction and every other day. Expression of LaccID and engineered receptors was verified at 96 h post-transduction by flow cytometry (Day 5). Cells co-expressing LaccID and the engineered receptor were sorted, expanded and maintained at a cell density of 0.5–1 million cells/ml until Day 14.

### Sample preparation for T cell proteomics

A375 melanoma tumor cells (ATCC CRL-1619) were cultured in complete DMEM (Gibco Cat# 11965-092) supplemented with 10% (v/v) fetal bovine serum (FBS, VWR Cat# 97068-085) and 1% penicillin-streptomycin (VWR Cat# 16777-164) at 37 °C under 5% CO_2_. Four million A375 cells were seeded in a 150 mm dish, and 24 hours later, when the tumor cell population reached approximately 10 million, 30 million CD8+ primary human T cells expressing LaccID and a constitutive engineered receptor (TCR or CAR) or T cells expressing only LaccID (no TCR or CAR) were added to the culture, achieving an effector to target ratio of 3:1. The T cells and A375 tumor targets were co-cultured at 37 °C with 5% CO2 for 24 hours. Subsequently, the cells were labeled with 250 μM BMP for 2 hours, and then quenched with quenchers (10 mM sodium ascorbate and 5 mM Trolox). The 40 million cells were pelleted and washed three times with DPBS containing quenchers. The cells were lysed in 1ml of RIPA lysis buffer (Millipore) containing protease inhibitor cocktail, radical quenchers, 10 mM sodium azide, and benzonase. 50 μl of the lysate were reserved for Western blotting, and the remainder was used for streptavidin enrichment following the protocol described in Cho et al.^7^ Streptavidin beads were washed twice with 1 ml of RIPA buffer and incubated with the cell lysate for 1 h at room temperature with rotation. The beads were then washed three times with 1 ml of RIPA buffer, followed by washes of 1 M KCl, 0.1 M Na_2_CO_2_ and 2 M urea in 10 mM Tris-HCl (pH 8.0). The beads were then washed with 1 ml of RIPA two more times. Finally, the beads were washed twice with 75 mM NaCl in 50 mM Tris-HCl (pH 8.0), frozen in 50 μl of the buffer, and stored until further processing.

### On-bead trypsin digestion of biotinylated proteins

Biotinylated proteins bound to streptavidin magnetic beads were subjected to four washes with 200 μl of 50 mM Tris-HCl (pH 7.5) buffer. After removing the final wash, the beads were incubated twice at room temperature (RT) in 80 μl of the digestion buffer – 2 M Urea, 50 nM Tris-HCl, 1 mM DTT, and 0.4 ug trypsin – while shaking at 1000 rpm. The first incubation lasted 1 hour, followed by the second incubation of 30 minutes. After each incubation, the supernatant was collected and transferred into a separate tube. The beads were then washed twice with 60 μl of 2M Urea/50 mM Tris-HCl buffer. The resulting washes were combined with the digestion supernatant. The pooled eluate of each sample was then spun down at 5000 x g for 30 sec to collect the supernatant. The samples were subsequently reduced with 4 mM DTT for 30 minutes at RT with shaking at 1000 rpm, followed by alkylation with 10 mM Iodoacetamide for 45 min in the dark at RT while shaking at 1000 rpm. Overnight digestion of the samples was performed by adding 0.5ug of trypsin to each sample. The following morning, the samples were acidified with neat formic acid (FA) to the final concentration of 1% FA (pH < 3.0).

Digested peptide samples were desalted using in-house packed C18 (3 M) StageTips. C18 StageTips were conditioned sequentially with 100 μl of 100% methanol (MeOH), 100 μl of 50% (vol/vol) acetonitrile (MeCN) with 0.1% (vol/vol) FA, and two washes of 100uL of 0.1% (v/v) FA. Acidified peptides were loaded onto the C18 StageTips and washed twice with 100 μl of 0.1% FA. The peptides were then eluted from the C18 resin using 50 μl of 50% MeCN/ 0.1% FA. The desalted peptide samples were snap-frozen and vacuum-centrifuged until completely dry.

### TMT labeling and fractionation

Desalted peptides were labeled with TMTpro reagents (Thermo Fisher Scientific). Each peptide sample was resuspended in 80 μL of 50 mM HEPES and labeled with 20 μl of the 25 ug/uL TMT reagents in MeCN. The samples were then incubated at RT for 1 hour while shaking at 1000 rpm. To quench the TMT-labeling reaction, 4 μL of 5% hydroxylamine was added to each sample, followed by a 15-minute incubation at RT with shaking. TMT-labeled samples were combined and vacuum-centrifuged to dry. The samples were then reconstituted in 200 μL of 0.1% FA and desalted on a C18 StageTips using the previously described protocol. The desalted TMT-labeled combined sample was then dried to completion.

The combined TMT-labeled peptide sample was fractionated by basic reverse phase (bRP) fractionation using an in-house packed SDB-RPS (3M) StageTip. The SDB-RPS StageTip was first conditioned sequentially with 100 μL of 100% MeOH, 100 μL 50% MeCN/0.1% FA, and two washes of 100 μL 0.1% FA. The peptide sample was reconstituted in 0.1% FA and loaded onto the SDB-RPS StageTip. Peptides were then eluted from the StageTip in eight fractions using 20 mM ammonium formate buffers with increasing (vol/vol) concentrations of MeCN (5%, 7.5%, 10%, 12.5%, 15%, 20%, 25%, and 45%). The eight fractions were then dried down to completion.

### Liquid chromatography and mass spectrometry

All peptide samples were separated and analyzed on an online liquid chromatography tandem mass spectrometry (LC-MS/MS) system, consisting of a Vanquish Neo UPHLC (Thermo Fisher Scientific) coupled to an Orbitrap Exploris 480 (Thermo Fisher Scientific). All peptide fractions were reconstituted in 9 μl of 3% MeCN/ 0.1% FA. 4 μl of each fraction was injected onto a microcapillary column (Picofrit with 10 µm tip opening / 75 µm diameter, New Objective, PF360-75-10-N-5), packed in-house with 30 cm of C18 silica material (1.5 µm ReproSil-Pur C18-AQ medium, Dr. Maisch GmbH, r119.aq) and heated to 50 °C using column heater sleeves (PhoenixST). Peptides were eluted into the Orbitrap Exploris 480 at a flow rate of 200 nL/min. The bRP fractions were run on a 110min-method, including a linear 84 min gradient from 94.6% solvent A (0.1% formic acid) to 27% solvent B (99.9% acetonitrile, 0.1% formic acid), followed by a linear 9 min gradient from 27% solvent B to 54% solvent B.

Mass spectrometry was conducted using a data-dependent acquisition mode, where MS1 spectra were measured with a resolution of 60,000, a normalized AGC target of 300%, and a mass range from 350 to 1800 m/z. MS2 spectra were acquired for the top 20 most abundant ions per cycle at a resolution of 45,000, an AGC target of 30%, an isolation window of 0.7 m/z and a normalized collision energy of 34. The dynamic exclusion time was set to 20 s, and the peptide match and isotope exclusion functions were enabled.

### Analysis of mass spectrometry data

Mass spectrometry data was processed using Spectrum Mill (proteomics.broadinstitute.org). Spectra within a precursor mass range of 600-6000 Da with a minimum MS1 signal-to-noise ratio of 25 were retained. Additionally, MS1 spectra within a retention time range of +/- 45 s, or within a precursor m/z tolerance of +/- 1.4 m/z were merged. MS/MS searching was performed against a human Uniprot database, released on September 02, 2021, supplemented with the sequence of LaccID to track its expression across samples. For searching, the full-mix function of the TMTpro modification with the fixed modification of carbamidomethylation on cysteine were used. Variable modifications included acetylation of the protein N-terminus, oxidation of methionine, cyclization to pyroglutamic acid, and N-term deamidation. Digestion parameters were set to “trypsin allow P” with an allowance of 4 missed cleavages. The matching tolerances were set with a minimum matched peak intensity of 30%, precursor and product mass tolerance of +/- 20 ppm, and a maximum ambiguous precursor charge of 3.

Peptide spectrum matches (PSMs) were validated with a maximum false discovery rate (FDR) threshold of 1.2% for precursor charges ranging from +2 to +6. A target protein score of 0 was applied during protein polishing autovalidation to further filter PSMs. TMTpro reporter ion intensities were corrected for isotopic impurities using the afRICA correction method in the Spectrum Mill protein/peptide summary module, which utilizes determinant calculations according to Cramer’s Rule. Protein quantification and statistical analysis were performed using the Proteomics Toolset for Integrative Data Analysis (Protigy, v1.0.7, Broad Institute, https://github.com/broadinstitute/protigy). Differential protein expression was evaluated using moderated t-tests, with P-values calculated to assess significance.

### Surface proteomic data analysis

To generate monoculture and coculture surfaceomes, a percent sequence coverage filter was applied with a 4% cutoff as it improved specificity of the datasets (**Supplementary Fig. 11**). Then, the filtered datasets were compared to the Omit LaccID condition, using a p-value cutoff of 0.01 and a false discovery rate (FDR) of 5% for monocultures and 10% for cocultures (**Supplementary Table 2** and **Supplementary Figs. 9f, 10b**). The FDR was calculated using a subset of false positive proteins expected not to be labeled by LaccID: nucleus and cytosol proteins that are not annotated with cell surface, Golgi, or membrane related terms (**Supplementary Table 4**). The cell surface annotation rate was calculated using a subset of proteins expected to be exposed on the cell surface, which are annotated in Uniprot^73^, Human Protein Atlas^74^, and GO Cellular Component^75, 76^ terms including: cell membrane, cell surface, secreted (secretory granule, synaptic vesicle), endosomes (early, late, recycling, membrane), lysosome, plasma membrane (basal, apical, lateral, external side, raft), extracellular regions (space, matrix, vesicle, exosome), immunological synapse, cilium, intercellular bridge, membrane raft, junctions (adherens, bicellular tight, anchoring, cell-cell), presynaptic membrane, early endosome, basement membrane, exocyst, ruffle membrane, and focal adhesion (**Supplementary Table 4**). Golgi and ER proteins used in **Fig. 5c** are listed in **Supplementary Table 4**.

To compare cocultures versus monocultures, filtered datasets of each were combined and filtered using a p-value cutoff of 0.01. Proteins were then ranked by Log_2_FC (coculture over monoculture). The GO cellular component and biological process of top 50 proteins and bottom 40 proteins were analyzed by g:Profiler^77^.

## Supporting information

Supplementary Figures 1-10, Genetic constructs list, Antibodies list, Legends of Supplementary Tables 1-4, Protein sequences of constructs

## Acknowledgements

We thank M. Sanchez (University of Cambridge) for initial assistance with directed evolution. This work was supported by the NIH (RC2DK129964 to A.Y.T., R01-DC006982 to L.L., and K99-DC021195 to C.N.M.), NSF (NeuroNex UTA20-000889 to A.Y.T. and 2014862 to M.H.E), Wu Tsai Neurosciences Institute Knight Initiative (to A.Y.T.), MCIN/AEI/doi: 10.13039/501100011033 grant no. I+D+I PID2022-142074OB-I00-RAINBOW (to M.A.), Comunidad de Madrid Atraccion de Talento Mod. 1 Project 2022-T1/BIO-238521/ECOCHEM. (to D.G.P.), and the Swiss National Science Foundation Postdoc Mobility fellowship (to D.H.). R.A.H.-L. holds a Career Award at the Scientific Interface from Burroughs Welcome Fund. A.Y.T. and R.A.H.-L. are Chan Zuckerberg Biohub – San Francisco Investigators.

## Data availability

The data associated with this study are available in the article and the Supplementary Information. The original mass spectra and the protein sequence database used for searches have deen deposited in the public proteomics repository MassIVE (http://massive.ucsd.edu) and are accessible at ftp://MSV000095684@massive.ucsd.edu. Additional data beyond that provided in the figures and Supplementary Information are available from the corresponding author on request.

