## Supplementary Figures 1-10, Genetic constructs list, Antibodies list, Legends of Supplementary Tables 1-4, Protein sequences of constructs for "Directed evolution of the multicopper oxidase laccase for cell surface proximity labeling and electron microscopy"

(Contents)

Supplementary Figures 1-10

Genetic constructs list

Antibodies list

Legends of Supplementary Tables 1-4

Protein sequences of constructs

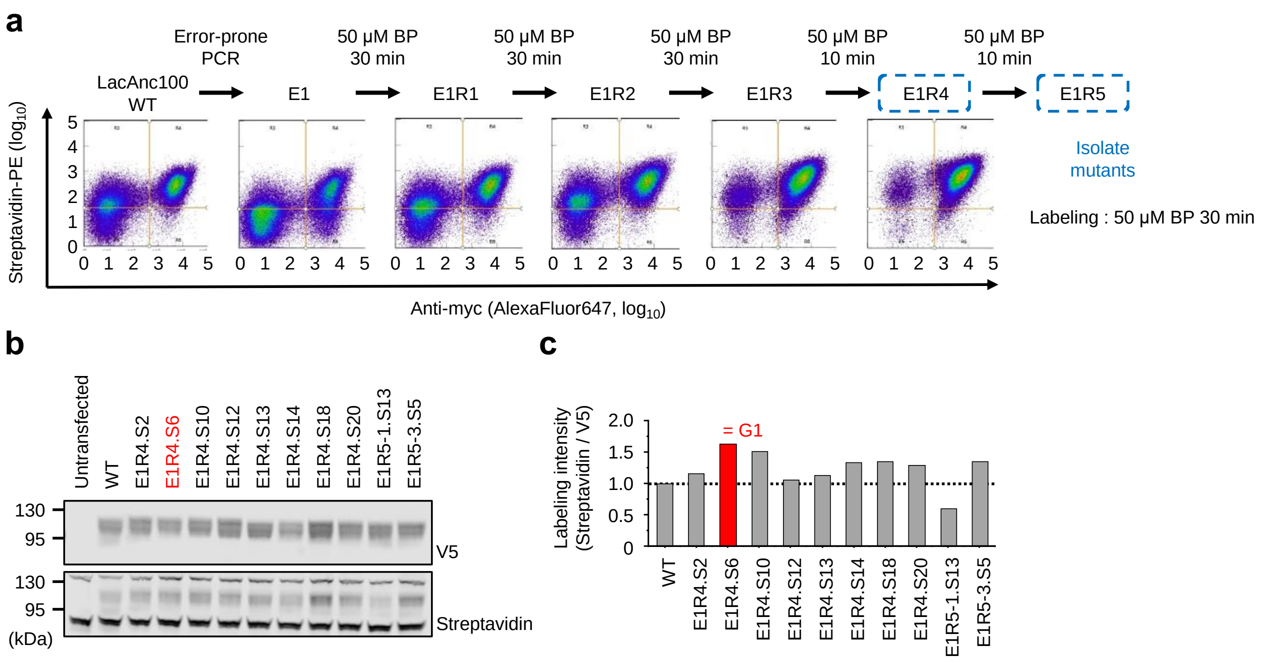


**Supplementary Figure 1. Directed evolution of laccase – first generation.** (**a**) FACS analysis showing progression of LacAnc100 library over 5 rounds of selection. E1 is the initial library derived from LacAnc100 template. E1R1 to E1R5 are the yeast populations recovered from Rounds 1-5. For this comparison, all samples were labeled with 50 μM BP for 30 min before streptavidin-PE + anti-myc staining. This experiment was performed once. (**b**) Streptavidin and anti-V5 blot analysis of first generation clones. LacAnc100 mutants were expressed on the surface of HEK293T cells and treated with 500 μM BP for 1 h in DMEM. (**c**) Quantification of data in **b**.

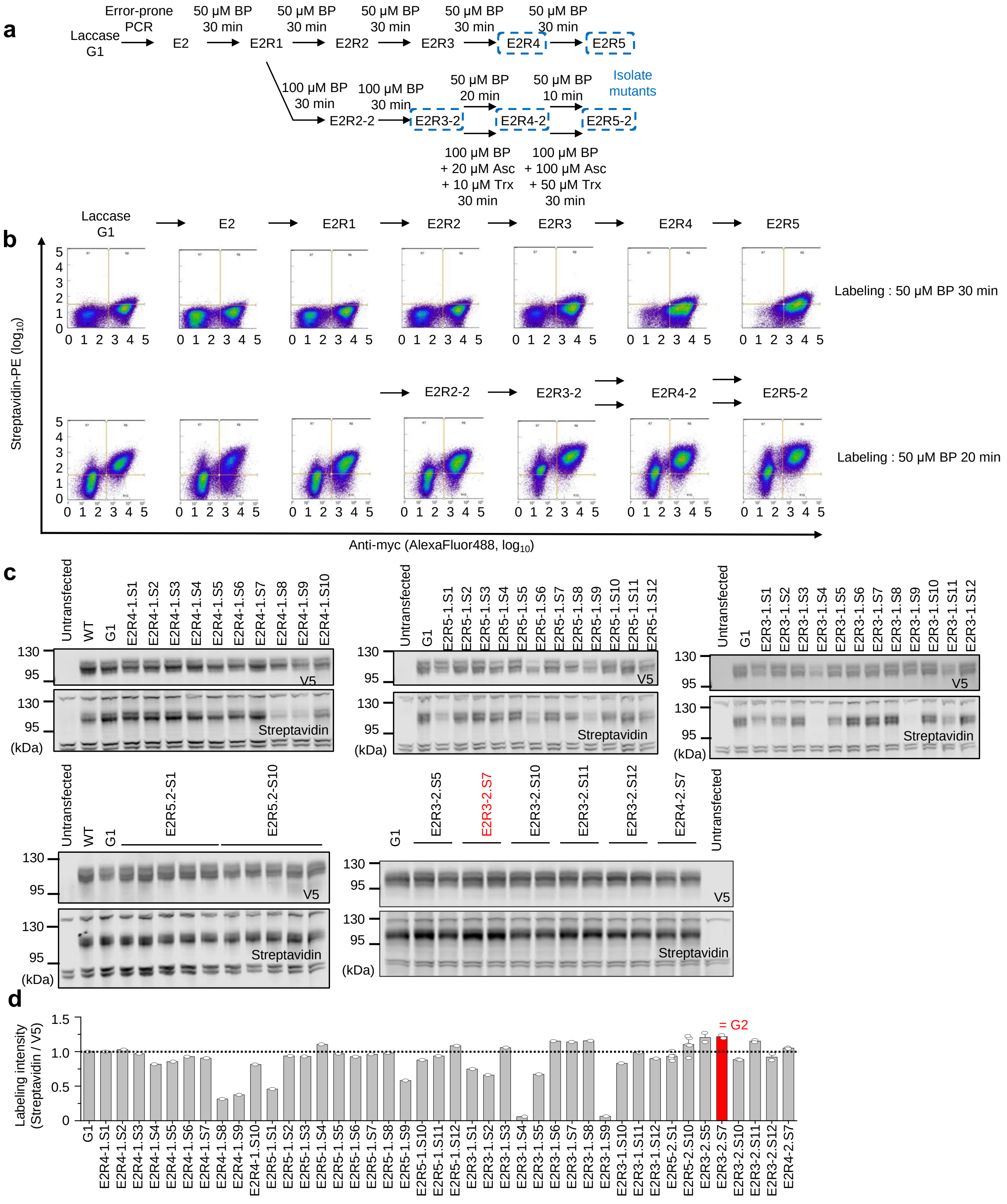


**Supplementary Figure 2. Directed evolution of laccase – second generation.** (**a**) Summary of selection conditions used in second generation evolution. E2 is the library derived from G1 template. (**b**) FACS analysis showing progression of library over 5 rounds of selection. For this comparison, all samples were labeled with 50 μM BP for 30 min (top) or 20 min (bottom) before streptavidin-PE + anti-myc staining. This experiment was performed once. (**c**) Streptavidin and anti-V5 blot analysis of second generation clones. LacAnc100 mutants were expressed on the surface of HEK293T cells and treated with 500 μM BP for 12 h in DMEM. (**d**) Quantification of data in **c**.

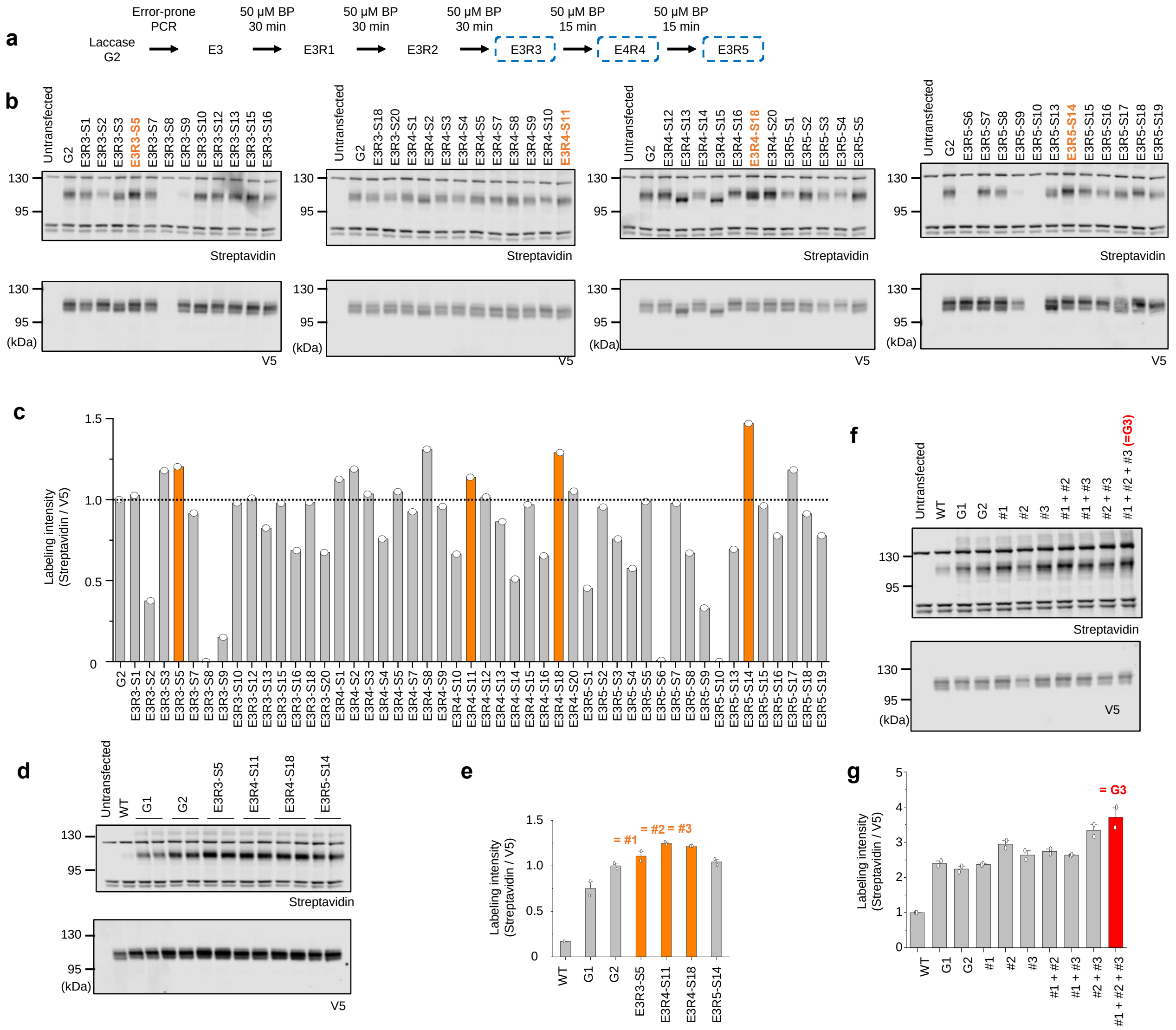

**Supplementary Figure 3. Directed evolution of laccase – third generation.** (**a**) Summary of selection conditions used in third generation directed evolution. E3 is the library derived from G2 template. (**b**) Streptavidin and anti-V5 blot analysis of third generation clones. LacAnc100 mutants were expressed on the surface of HEK293T cells and treated with 500 μM BP for 12 h in DMEM. (**c**) Quantification of data in **b**. (**d**) Streptavidin and anti-V5 blot analysis of four selected clones (colored orange in **b**, **c**). Individual clones were expressed on the surface of HEK293T cells and treated with 500 μM BP for 12 h in DMEM. (**e**) Quantification of data in **d**. (**f**) Streptavidin blotting analysis of four clones that have combined mutations. Individual clones were expressed on the surface of HEK293T cells and treated with 500 μM BP in DMEM. (**g**) Quantification of data in **f**.

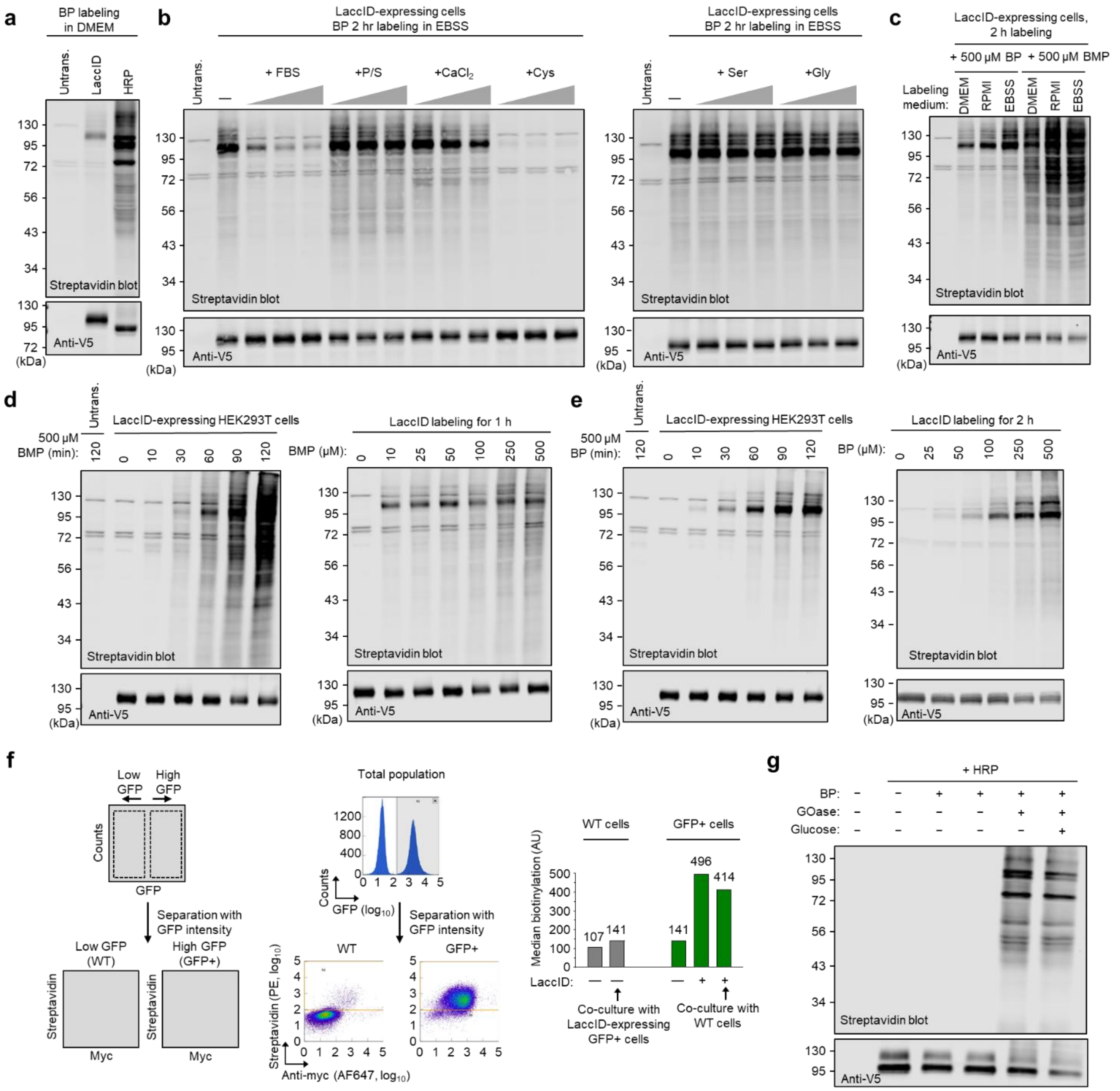


**Supplementary Figure 4. Additional characterization of LaccID, related to Figure 2.** (**a**) LaccID + BP 2 h in DMEM is still not as efficient as HRP labeling for 1 min. (**b**) LaccID labeling in EBSS in the presence of 5, 10, 15% FBS, 0.5, 1, 1.5% penicillin-streptomycin, 50, 100, 150 mM $CaCl_2$, and 0.03, 0.06, 0.09 g/L cysteine, glycine, or serine. (**c**) LaccID labeling in different cell media. HEK293T cells expressing surface LaccID were labeled for 2 h with 500 μM BP or BMP. (**d**) LaccID labeling time course and substrate titration for BMP. (**e**) LaccID labeling time course and substrate titration for BP. (**f**) Characterization of trans-labeling extent with BMP using K562 cells in suspension. K562 cells expressing both cytosolic GFP and cell surface LaccID (GFP+ LaccID+) were co-cultured in a 1:1 ratio with non-expressing K562 cells. The total cell density was 9 x $10^9$ cells/mL. After labeling with 500 μM BMP in RPMI for 2 h, co-cultures were stained with streptavidin-PE and anti-myc antibody and analyzed by FACS. LaccID+ and non-expressing (wild-type) cells were analyzed separately by first gating on GFP intensity. Middle: representative GFP histograms and streptavidin vs. myc 2D scatter plots are shown. Right: quantification of wild-type cells and GFP+ cells showing a small degree of trains-labeling on the former when co-cultured with LaccID+ cells. (**g**) HRP can use $H_2O_2$ generated by Glucose oxidase (GOase). HRP showed increased labeling when glucose oxidase was added, likely due to the generation of $H_2O_2$ from exogenously-added glucose or residual glucose from cell culture media.

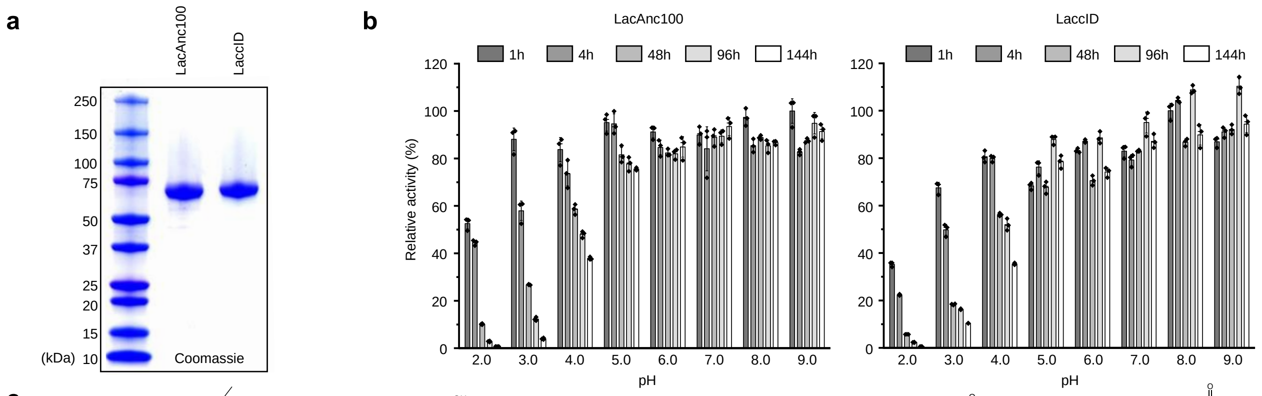


c

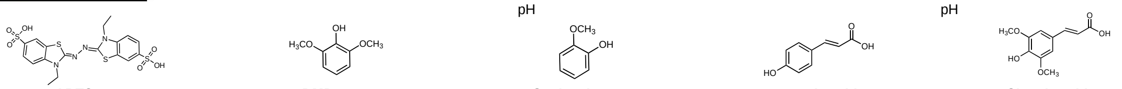

| Variant | ABTS ITN (μmol product·μmol$^{-1}$·min$^{-1}$) | ABTS Fold increase | DMP ITN (μmol product·μmol$^{-1}$·min$^{-1}$) | DMP Fold increase | Guaiacol ITN (μmol product·μmol$^{-1}$·min$^{-1}$) | Guaiacol Fold increase | *p*-coumaric acid ITN (μmol product·μmol$^{-1}$·min$^{-1}$) | *p*-coumaric acid Fold increase | Sinapic acid ITN (μmol product·μmol$^{-1}$·min$^{-1}$) | Sinapic acid Fold increase |
|---|---|---|---|---|---|---|---|---|---|---|
| LacAnc100 | 339 ± 10 | - | 200 ± 3 | - | 48 ± 18 | - | 18 ± 0.4 | - | 339 ± 29 | - |
| LaccID | 1,899 ± 24 | 5.6 | 1,744 ± 48 | 8.7 | 1,000 ± 5 | 20.8 | 20 ± 0.1 | 1.1 | 2,823 ± 89 | 8.3 |

d

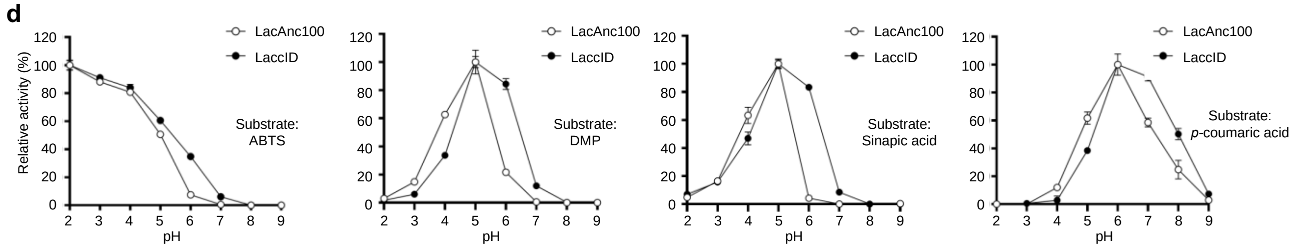


e

Kinetic constants for ABTS at pH 6.0.

| Kinetic constants | LacAnc100 | LaccID |
|---|---|---|
| $K_m$ (mM) | 0.032 ± 0.001 | 0.083 ± 0.002 |
| $k_{cat}$ ($s^{-1}$) | 173 ± 1 | 498 ± 5 |
| $k_{cat}/K_m$ ($mM^{-1}$. $s^{-1}$) | 5,272 ± 117 | 5,992 ± 111 |

Kinetic constants for DMP at pH 6.0.

| Kinetic constants | LacAnc100 | LaccID |
|---|---|---|
| $K_m$ (mM) | 0.099 ± 0.007 | 0.20 ± 0.01 |
| $k_{cat}$ ($s^{-1}$) | 104 ± 2 | 310 ± 5 |
| $k_{cat}/K_m$ ($mM^{-1}$. $s^{-1}$) | 1,051 ± 57 | 1,491 ± 50 |

Kinetic constants for Sinapic acid at pH 6.0.

| Kinetic constants | LacAnc100 | LaccID |
|---|---|---|
| $K_m$ (mM) | 0.038 ± 0.006 | 0.1 ± 0.02 |
| $k_{cat}$ ($s^{-1}$) | 86 ± 4 | 177 ± 14 |
| $k_{cat}/K_m$ ($mM^{-1}$. $s^{-1}$) | 2,257 ± 288 | 1,743 ± 171 |

f

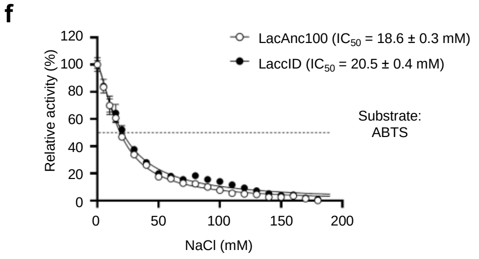


**Supplementary Figure 5. Characterization of LaccID in vitro, related to Figure 2.** (**a**) Coomassie blue staining of a 4-20% SDS-PAGE gel showing electrophoresis results of purified LacAnc100 and LaccID by immobilized metal affinity chromatography. (**b**) In vitro pH stability profiles of LacAnc100 and LaccID. (**c**) Initial turnover numbers (ITN) with different substrates for LacAnc100 and LaccID. Measurements were performed in PBS at pH 7.4. (**d**) In vitro pH activity profiles of LacAnc100 and LaccID. (**e**) Kinetic constants of LacAnc100 and LaccID for ABTS, DMP, and Sinapic acid at pH 6.0. (**f**) Halide inhibition of LacAnc100 and LaccID. $IC_{50}$, half-maximal inhibitory concentration of NaCl.

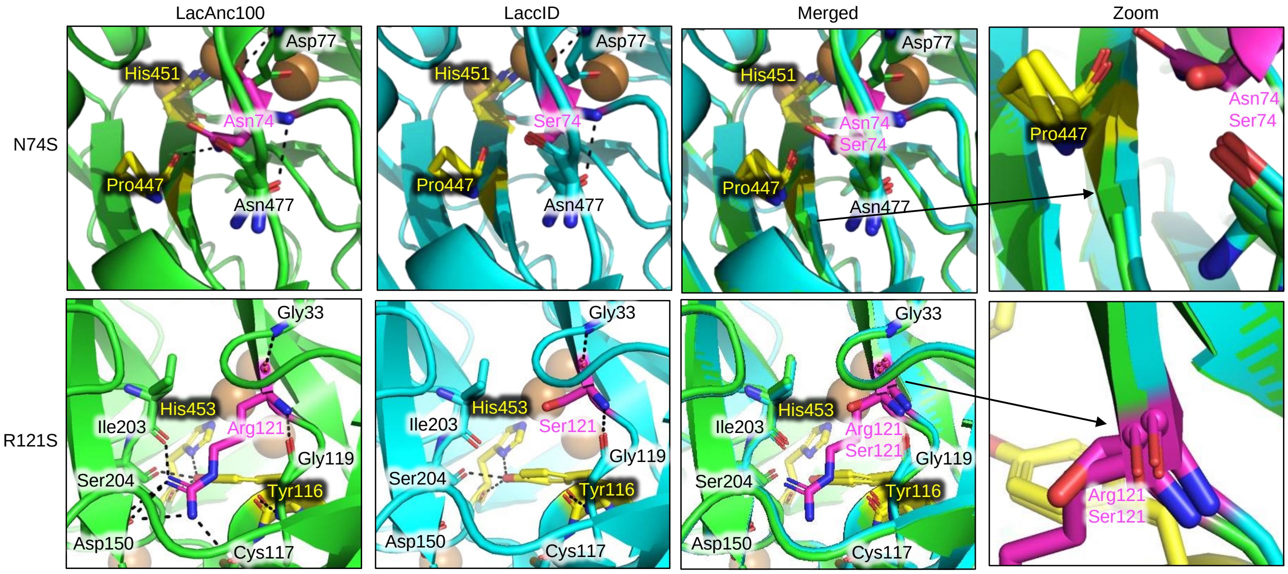


**Supplementary Figure 6. Structural analysis on LaccID mutations.** Structural changes in LaccID driven by the N74S and R121S mutations can be predicted by AlphaFold3. In all 5 predicted structures, structural changes were observed around two mutations when compared to LacAnc100. The N74S mutation in LaccID disrupts the interaction with Pro447, resulting in a shift in the beta-strand containing the copper-coordinating His451 residue. The R121S mutation shifts another beta-strand and also breaks an interaction with Tyr116, which interacts with another copper-coordinating residue His453. It is worth noting that His451 and His453 both are part of the HCH motif, which is crucial for LaccID activity (**Fig. 2e**). Histidines and their interacting amino acids are shown in yellow and copper atoms in orange.

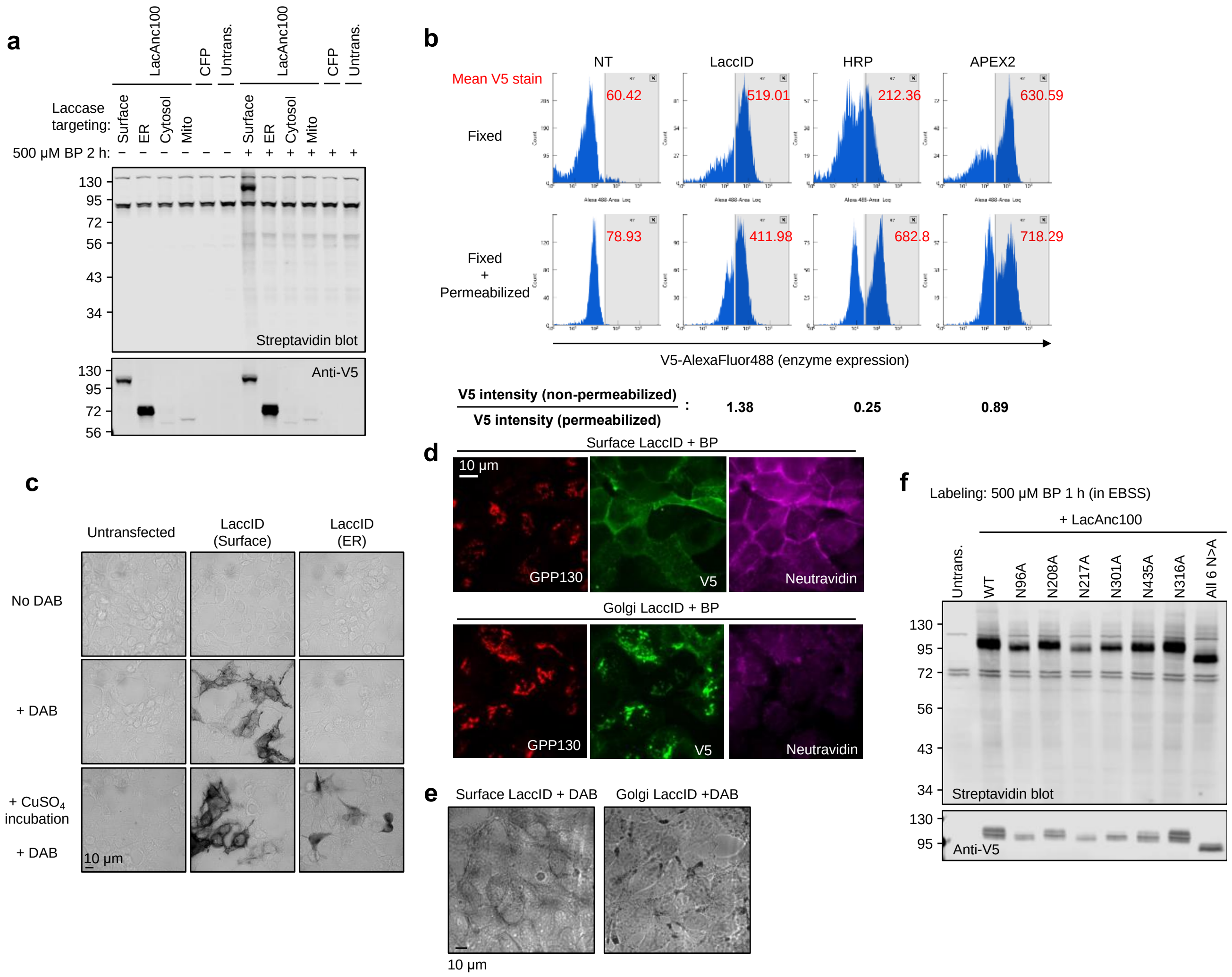


**Supplementary Figure 7. LaccID is specifically active on the cell surface, related to Figure 3.** (**a**) LacAnc100 is only active on the cell surface. HEK293T cells expressing LacAnc100 targeted to the cell surface, ER lumen, cytosol, or mitochondria were labeled for 2 h with 500 μM BP. (**b**) LaccID shows better surface targeting compared to HRP and APEX2. HEK293T cells expressing surface-targeted enzymes were fixed and stained with anti-V5-AF488, with or without permeabilization. HRP and APEX2 show increase in anti-V5 intensity upon cell permeabilization, indicating that more enzymes are present in intracellular compartments. (**c**) DAB staining shows that ER-LaccID could be activated by adding exogenous $CuSO_4$. (**d**) Golgi-targeted LaccID does not exhibit BP labeling in HEK293T cells. After labeling for 2 h with 500 μM BP, cells were fixed and stained with anti-V5 antibody to detect expression, neutravidin-AF647 to detect biotinylated proteins, and anti-GPP130 antibody to stain the Golgi. (**e**) Golgi-targeted LaccID showed DAB staining in fixed cells. HEK293T cells expressing LaccID targeted to the cell surface or the Golgi apparatus were fixed and then incubated with DAB for 1h. (**f**) Mutants of LacAnc100 to remove N-glycosylation are still active.

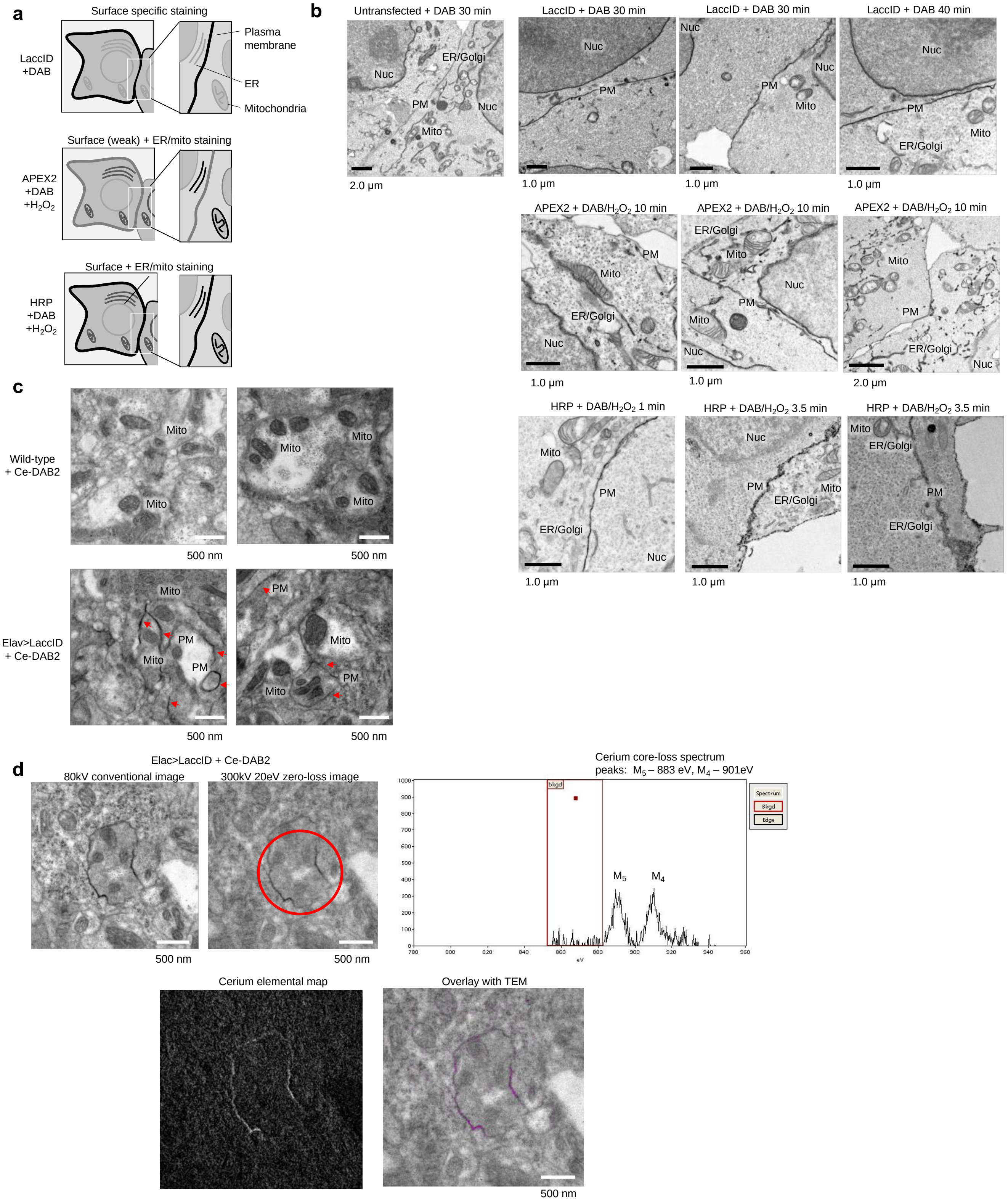


**Supplementary Figure 8. Application in EM, related to Figure 4.** (**a**) Expected EM staining patterns for surface-targeted LaccID, HRP, and APEX2. (**b**) EM imaging of HEK293T cells expressing cell surface-targeted enzymes. Cells were fixed using 2% glutaraldehyde, and incubated with DAB for 30 min, followed by post-fixation with 2% reduced osmium tetroxide. PM, plasma membrane. Nuc, nucleus. (**c**)

Additional fields of view of EM images of DAB-stained fly ventral nerve cord (VNC), related to **Fig. 4g**. (**d**) Electron Energy Loss Spectroscopy (EELS) confirms the presence of cerium in the dark DAB stain. Left: EM image of Ce-DAB2-stained fly VNC. Right: EELS spectra collected from the region marked in left panel in red. Characteristic peaks at 883 (M5) and 901 (M4) eV confirm the presence of cerium.

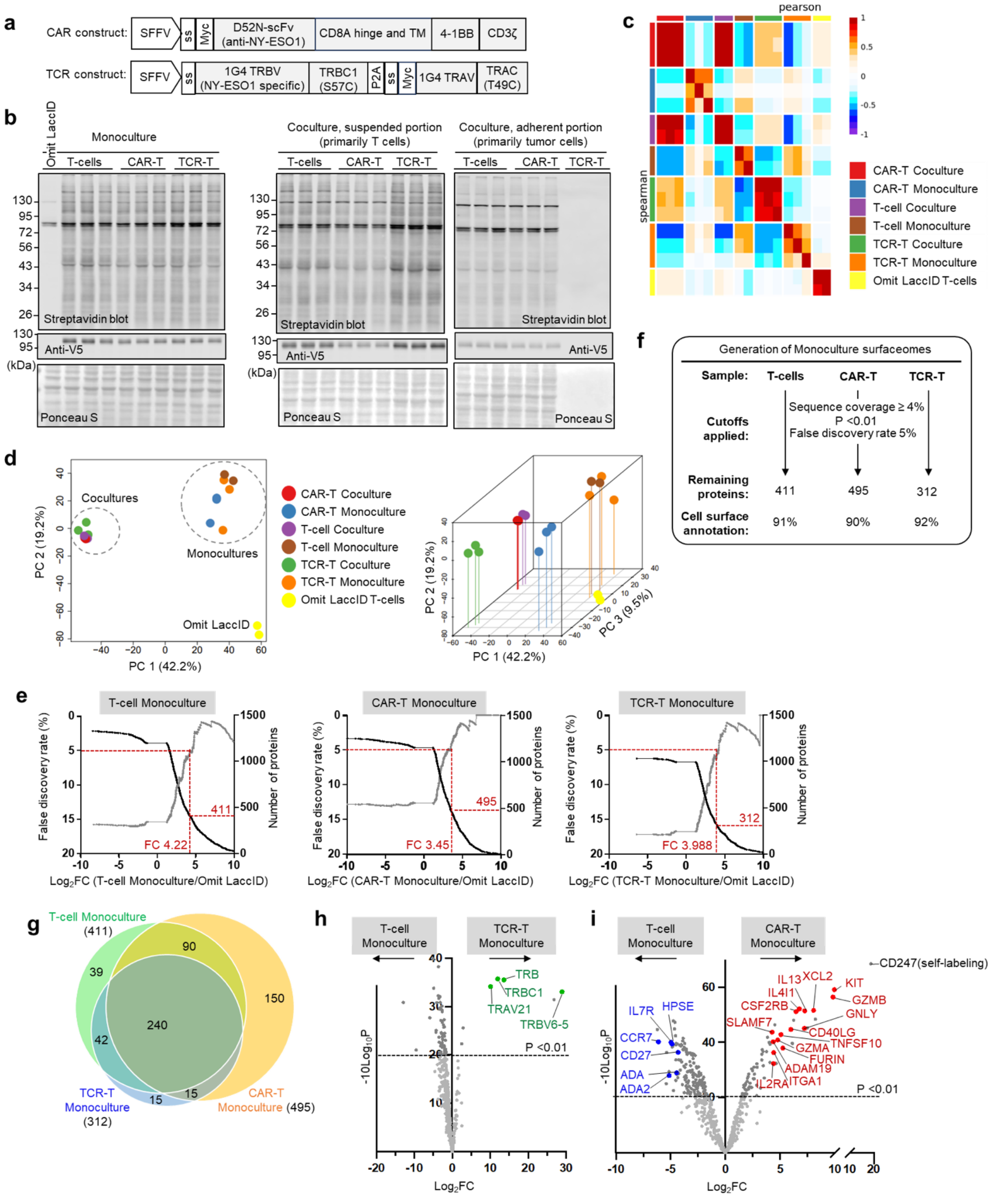


**Supplementary Figure 9. Additional data related to LaccID mapping of T cell surface proteomes.** (**a**) Constructs used for CAR and TCR expression in CD8+ T-cells. (**b**) Streptavidin blot analysis of BMP-labeled proteomic samples from **Fig. 5b**. Staining is largely absent in TCR-T cocultures (adherent portion) because tumor cells have been killed by the TCR-T cells. (**c**) Correlation between biological replicates. (**d**) Principal component analysis of proteomic samples. (**e**) Procedure for generation of monoculture surfaceomes. (**f**) False discovery rate (FDR) and proteome size as a function of Log2 fold cutoff. 5% FDR was used to select cutoffs for all three monoculture surfaceomes. (**g**) Venn diagram comparing content of three monoculture surfaceomes. (**h**) Volcano plots comparing TCR-T monoculture to T-cell monoculture surfaceomes. The surfaceome of TCR-T cells is very similar to that of T-cells and the primary differences are exogenously expressed TCR components, specifically IG4 TRBV (TRB and TRBV6-5), TRBC1, 1G4 TRAV (TRAV21), and TRBC1. The expressed TCR components are colored green. (**i**) Volcano plots comparing CAR-T monoculture to T-cell monoculture surfaceomes. Proteins with prior literature connections to upregulation upon T-cell activation are colored red. Proteins with prior literature connections to downregulation upon T-cell activation, or those known to improve CAR-T cell efficacy when overexpressed or upregulated, are colored blue.

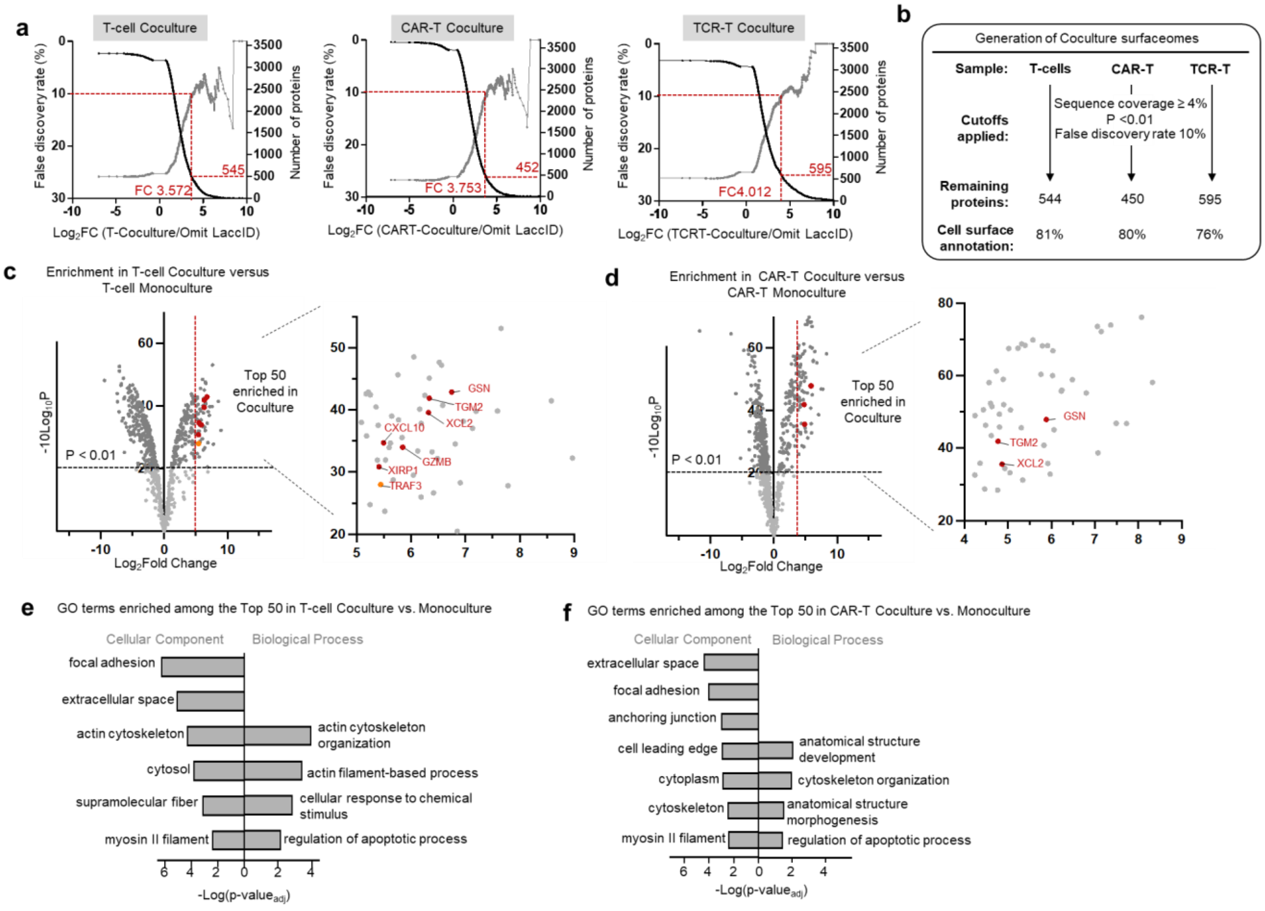


**Supplementary Figure 10. Additional analyses of LaccID-labeled T-cell: tumor cell cocultures.** (**a**) False discovery rate (FDR) and proteome size as a function of Log2 fold cutoff. 10% FDR was used to select cutoffs for all three coculture surfaceomes. (**b**) Procedure for generation of surfaceomes enriched in cocultures versus monocultures. (**c**) Volcano plots comparing T-cell coculture to T-cell monoculture. Proteins with prior literature connection to T-cell activation are colored red. (**d**) Volcano plot comparing CAR-T coculture to CAR-T monoculture. Proteins with prior literature connection to T-cell activation are colored red. (**e**) Gene ontology terms enriched among the top 50 in T-cell coculture dataset compared to T-cell monoculture. (**f**) Gene ontology terms enriched among the top 50 in CAR-T coculture dataset compared to CAR-T monoculture.

### Genetic constructs

Table of genetic constructs used in this study

| Name | Promoter | Details | Used in |
|---|---|---|---|
| pDisplay_IgK signal sequence-HA-V5-Enzyme-CD4 TM | CMV | Target: Cell surface<br>Enzymes:<br>LacAnc100, ChuB, OB1, KyLO, G1, G2, LaccID, HRP, APEX2, LaccID(mut), LacAnc100 (N-to-A mut)<br>IgK: mouse IgK signal peptide, METDTLLLWVLLLWVPGSTGD<br>HA: YPYDVPDYA<br>V5: GKPIPNPLLGLDST<br>CD4 TM: human CD4 transmembrane domain, 202-427aa | Expression of enzyme in HEK293T cell surface |
| pDisplay_IgK signal sequence-HA-V5-Enzyme-KDEL | CMV | Target: ER lumen<br>Enzymes:<br>LacAnc100, LaccID, HRP, APEX2 | Expression of enzyme in HEK293T cell endoplasmic reticulum |
| pDisplay_HA-V5-LacAnc100-NES | CMV | Target: Cytosol<br>NES: LQLPPLERLTLD | Expression of LacAnc100 in HEK293T cell cytosol |
| pDisplay_MTS-HA-V5-LacAnc100 | CMV | Target: Mitochondrial matrix<br>MTS: human COX4I1 1-24aa, MLATRVFSLVGKRAISTSVCVRAH | Expression of LacAnc100 in HEK293T cell mitochondrial lumen |
| pTRE3G_IgK signal sequence-HA-V5-Enzyme-CD4 TM | TRE3G | Enzymes:<br>LacAnc100, G1, G2, LaccID, HRP, APEX2 | Stable expression of enzyme in HEK293T cell surface |
| pTRE3G_ManII-LaccID-V5 | TRE3G | Target: Golgi lumen<br>ManII: Mouse Man2a1 transmembrane domain, 1-97aa | Stable expression of enzyme in HEK293T cell surface |
| pCTCON2_Aga2p-myc-LaccID | GAL1 | LaccID and its variants | For yeast display of LaccID and its variants |
| pJRoC30_LaccID_8xHis | GAL1 | LaccID and LacAnc100 | For purification of recombinant LaccID and its variants |
| pUASt_10xUAS-wgSS-HA-V5-LaccID-CD2 | UAS | | For expression of LaccID in *Drosophila* |

### Materials

Table of antibodies used in this study.

| Antibody | Source | Vendor | Catalog Number | Dilution(s) |
|---|---|---|---|---|
| Anti-V5 | Mouse | Thermo Fischer Scientific | R96025 | WB: 1:5000; IF: 1:3000 |
| Anti-myc | Chicken | Exalpha Biologicals, Inc. | ACMYC | Flow: 1:400 |
| Streptavidin-phycoerythin (PE) | - | Jackson ImmunoResearch | 0.16110084 | Flow: 1:200 |
| Anti-N-Cadherin | Rabbit | Thermo Fischer Scientific | PA5-19486 | WB: 1:5000; IF: 1:2000 |
| Anti-Calreticulin | Rabbit | Thermo Fischer Scientific | PA3900 | WB: 1:5000; IF: 1:2000 |
| Anti-mouse-HRP | Goat | BioRad | 1706516 | WB: 1:3000 |
| Anti-rabbit-HRP | Goat | BioRad | 1706515 | WB: 1:3000 |
| DAPI | - | Enzo Life Sciences | AP402-0010 | IF: 1 μg/mL final |
| Anti-mouse-AlexaFluor488 | Goat | Invitrogen | A11029 | IF: 1:1000 |
| Anti-rabbit-AlexaFluor568 | Goat | Invitrogen | A11036 | IF: 1:1000 |
| Anti-chicken-AlexaFluor488 | Goat | Invitrogen | A11039 | Flow: 1:200 |
| Anti-chicken-AlexaFluor647 | Goat | Invitrogen | A21449 | Flow: 1:200 |
| Neutravidin-AlexaFluor647 | - | Thermo Fischer Scientific | A2666; A20006 | IF: 1:1000 |
| Anti-mouse-IRDye 800CW | Goat | LI-COR Biosciences | 92632210 | WB: 1:5000 |
| Anti-mouse-IRDye 680CW | Goat | LI-COR Biosciences | 92668070 | WB: 1:5000 |
| Anti-rabbit-IRDye 800CW | Goat | LI-COR Biosciences | 92632211 | WB: 1:5000 |
| Anti-rabbit-IRDye 680CW | Goat | LI-COR Biosciences | 92568071 | WB: 1:5000 |

| | | | | |
|---|---|---|---|---|
| Streptavidin-IRDye 680CW | - | LI-COR Biosciences | 92568079 | WB: 1:5000 |
| Anti-myc-AlexaFluor647 | Mouse | Santa Cruz Biotechnology | sc-40 AF647 | Flow: 1:200 |
| Anti-V5-iFluor488 | Mouse | Genscript | A01803-100 | Flow: 1:500 |

**Supplementary Table 1**. **Tab 1:** Mutations found in individual clones in first generation evolution. Clone in red was named G1 and used for second generation evolution. **Tab 2**: Mutations found in individual clones in second generation evolution. Red one was named G2 and used for third generation evolution. **Tab 3.** Mutations found in individual clones in third generation evolution. Red clone was named LaccID.

**Supplementary Table 2**. T cell surfaceomes labeled by surface-targeted LaccID. **Tab 1**: T-cell monoculture surfaceomes. **Tab 2**: CAR-T monoculture surfaceomes. **Tab 3**: TCR-T monoculture surfaceomes. **Tab 4**: T-cell coculture surfaceomes. **Tab 5**: CAR-T coculture surfaceomes. **Tab 6**: TCR-T coculture surfaceomes.

**Supplementary Table 3**. Enriched proteins in coculture vs. monoculture. **Tab 1**: Top 50 proteins in TCR-T coculture vs. monoculture. **Tab 2**: Bottom 40 proteins in TCR-T coculture vs. monoculture. **Tab 3**: Top 50 proteins in T-cell coculture vs. monoculture. **Tab 4**: Top 50 proteins in CAR-T coculture vs. monoculture.

**Supplementary Table 4**. **Tab 1**: Unfiltered proteomic data. **Tab 2**: False positives used to filter the proteomic data (described in **Methods**). **Tab 3**: Surface annotated protein list used for surface annotation rate calculation. **Tab 4**: Golgi-resident protein list used in **Fig. 5c**. **Tab 5**: ER-resident protein list used in **Fig. 5c**.

### Protein sequences

**HA-V5-LacAnc100-CD4 TM (Surface LacAnc100)**

METDTLLLWVLLLWVPGSTGDYPYDVPDYAGAQPARSGGGKPIPNPLLGLDSTASAIGPVADLHIVNA
DISPDGFTRSAVLAGGTFPGPLITGNKGDNFQINVINQLTDTTMLRSTSIHWHGFFQHGTNWADGPAFV
TQCPIAPGNSFLYNFTVPDQAGTFWYHSHLSTQYCDGLRGPFVIYDPNDPHRSLYDVDDESTVITLAD
WYHTPAPQLAAAPPVPDSTLINGKGRYAGGPTSPLAVINVEQGKRYRFRLISISCDPNFTFSIDGHNMTI
IEADGVNTQPLTVDSIQIFAGQRYSFVLNANQPVDNYWIRANPNIGTTGFAGGINSAILRYVGAPEADPT
TTQTTSTNPLNETNLHPLENPGAPGNPTPGGADVNINLNIGFSSTGQFTINGVSFIPPTVPVLLQILSGAR
TAQDLLPSGSVYTLPPNKVIEISMPGGASAAGGPHPFHLHGHTFDVVRSAGSSTYNYVNPVRRDVVSI
GTAGDNVTIRFTTDNPGPWFLHCHIDWHLEAGLAVVFAEDTPDIASANPVPAAWDDLCPIYDALSPDD
QPRLQVDLDFQKASSIVYKKEGEQVEFSFPLAFTVEKLTGSGELWWQAERASSSKSWITFDLKNKEVS
VKRVTQDPKLQMGKKLPLHLTLPQALPQYAGSGNLTLALEAKTGKLHQEVNLVVMRATQLQKNLTCEV
WGPTSPKLMLSLKLENKEAKVSKREKAVWVLNPEAGMWQCLLSDSGQVLLESNIKVLPTWSTPVQPM
ALIVLGGVAGLLLFIGLGIFFCVRCRHRRR

Red: IgK signal peptide, Orange: HA epitope tag, Green: V5 epitope tag, Blue: LacAnc100, Purple: CD4 transmembrane domain

**HA-V5-ChuB-CD4 TM (Surface Chu-B)**

METDTLLLWVLLLWVPGSTGDYPYDVPDYAGAQPARSGGGKPIPNPLLGLDSTASSIGPVADLTISNGA
VSPDGFSRQAILVNDVFPSPLITGNKGDRFQLNVIDNMTNHTMLKSTSIHWHGFFQHGTNWADGPAFV
NQCPISTGHAFLYDFQVPDQAGTFWYHSHLSTQYCDGLRGPIVVYDPQDPHKSLYDVDDDSTVITLAD
WYHLAAKVGPAAPTADATLINGLGRSINTLNADLAVITVTKGKRYRFRLVSLSCDPNYTFSIDGHSLTVIE
ADGVNLKPQTVDSIQIFPAQRYSFVLNADQDVDNYWIRALPNSGTRNFDGGVNSAILRYEGAAPVEPTT
TQTPSTQPLVESALTTLEGTAAPGNPTPGGVDLALNMAFGFAGGRFTINGASFTPPTVPVLLQILSGAQ

SAQDLLPSGSVYSLPANADIEISLPATSAAPGFPHPIHLHGHTFAVVRSAGSSTYNYANPVYRDVVNTG
SPGDNVTIRFRTDNPGPWFLHCHIDEHLEAGFTVVMAEDIPDVAATNPVPQAWSDLCPTYDALSPDDQ
PRLQVDLDFQKASSIVYKKEGEQVEFSFPLAFTVEKLTGSGELWWQAERASSSKSWITFDLKNKEVSV
KRVTQDPKLQMGKKLPLHLTLPQALPQYAGSGNLTLALEAKTGKLHQEVNLVVMRATQLQKNLTCEV
WGPTSPKLMLSLKLENKEAKVSKREKAVWVLNPEAGMWQCLLSDSGQVLLESNIKVLPTWSTPVQPM
ALIVLGGVAGLLLFIGLGIFFCVRCRHRRR

Red: IgK signal peptide, Orange: HA epitope tag, Green: V5 epitope tag, Blue: Chu-B laccase, Purple: CD4 transmembrane domain

**HA-V5-OB-1-CD4 TM (Surface OB-1)**

METDTLLLWVLLLWVPGSTGDYPYDVPDYAGAQPARSGGGKPIPNPLLGLDSTASSIGPVADLTISNGA
VSPDGFSRQAILVNDVFPSPLITGNKGDRFQLNVIDNMTNHTMLKSTSIHWHGFFQHGTNWADGPAFV
NQCPISTGHAFLYDFQVPDQAGTFWYHSHLSTQYCDGLRGPIVVYDPQDPHKSLYDVDDDSTVITLAD
WYHLAAKVGPAAPTADATLINGLGRAAGGDADAALAVFNVTQGSRYRFRLVSLSCDPNFNFTIQDHNM
TIIEVDGVNVEPVTVDSIQIFAGQRYSFVLTADQDIGNYWIQAVPNTGTVTTDGGVNSAILRYDTADPIEP
DAADPTSSIPLVETDLVPLENTAAPGNPTPGGVDLAMNLEFDFNGTWFFINGEPFVPPSVPVLLQIMSG
AQSAADLLPSGSVYTLPANSTIEISFPMNTTAAPGAPHPFHLHGHTFYVVRSAGSTEYNYVNPPQRDTV
STGTDGDNVTIRFTTNNPGPWFLHCHIDFHLDAGFAVVMAEDTPDTKAANPVPQAWSDLCPIYDALDP
SDLPRLQVDLDFQKASSIVYKKEGEQVEFSFPLAFTVEKLTGSGELWWQAERASSSKSWITFDLKNKE
VSVKRVTQDPKLQMGKKLPLHLTLPQALPQYAGSGNLTLALEAKTGKLHQEVNLVVMRATQLQKNLTC
EVWGPTSPKLMLSLKLENKEAKVSKREKAVWVLNPEAGMWQCLLSDSGQVLLESNIKVLPTWSTPVQ
PMALIVLGGVAGLLLFIGLGIFFCVRCRHRRR

Red: IgK signal peptide, Orange: HA epitope tag, Green: V5 epitope tag, Blue: OB-1 laccase, Purple: CD4 transmembrane domain

**HA-V5-KyLO-CD4 TM (Surface KyLO)**

METDTLLLWVLLLWVPGSTGDYPYDVPDYAGAQPARSGGGKPIPNPLLGLDSTASQQICNTPSNRAC
WTDGYDINTDYEVDSPDTGVVRPYTLTLTEVDNWTGPDGVVKEKVMLVNNSIIGPTIFADWGDTIQVTV
INNLGTNGTSIHWHGLHQKGTNLHDGVSGITECPIPPKGGRKVYRFKAQQYGTSWYHSHFSAQYGNG
VVGAIQINGPASLPYDTDLGVFPISDYYYSSADKLVELTKNSGAPFSDNVLFNGTAKHPETGEGEYANV
TLTPGRRHRLRLINTSVENHFQVSLVNHTMTIIAADMVPVNAMTVDSLFLGVGQRYDVVIEANRTPGNY
WFNVTFGGGLLCGGSRSPYPAAIFHYAGAPGGPPTDEGKAPVDHNCLDLPNLKPVVARDVPLSGLAK
RPDNTLDVTLDTMGTPLFVWKVNGSAINIDWDRPVVDYVLTQNTSFPPGHNIVEVNGADQWSYWLIEN
DPGAPFTVPHPMHLHGHDFYVLGRSPDEPPASKERHVFDPARDAGLLSGANPVRRDVTMLPAFGWV
VLAFRADNPGAWLFHCHIAWHVSGGLGVVYLERADDLRGAVSEADADDFDRLCADWRRYWPTNPNP
KSDSGLKRRWVEEDKWLVKAPRLQVDLDFQKASSIVYKKEGEQVEFSFPLAFTVEKLTGSGELWWQA
ERASSSKSWITFDLKNKEVSVKRVTQDPKLQMGKKLPLHLTLPQALPQYAGSGNLTLALEAKTGKLHQ
EVNLVVMRATQLQKNLTCEVWGPTSPKLMLSLKLENKEAKVSKREKAVWVLNPEAGMWQCLLSDSG
QVLLESNIKVLPTWSTPVQPMALIVLGGVAGLLLFIGLGIFFCVRCRHRRR

Red: IgK signal peptide, Orange: HA epitope tag, Green: V5 epitope tag, Blue: KyLO laccase, Purple: CD4 transmembrane domain

**HA-V5-LaccID-CD4 TM (Surface LaccID)**

METDTLLLWVLLLWVPGSTGDYPYDVPDYAGAQPARSGGGKPIPNPLLGLDSTASAIGPVADLHIANA
DISPDGFTRSAVLAGGTFPGPLITGNKGDNFRINVINQLTDTTMLRSTSIHWHGFFQHGTSWADGPAFV
TQCPIAPGNSFLYNFTVPDQAGTFWYHSHLSTQYCDGLSGPFVVYDPNDPHRSLYDVDDESTVITLAD
WYHTPAPQLAAAPPVPDSTLINGKGRYAGGPTSPLAVINVEQGKRYRFRLISISCDPNFTFSIDGHNMTI
IEADGVNTQPLTVDSIQIFAGQRYSFVLNANQPVGNYWIRANPNIGTTGFAGGINSAILRYVGAPEADPT

TTQTTSTNPVNETNLHPLENPGAPGNPTPGGADVNINLNIGFSSTGQFTINGVSFIPPTVPVLLQILSGA
RTAQDLLPSGSVYTLPPNKVIEISMPGGASAAGGPHPFHLHGHTFDVVRSAGSSTYNYVNPVRRDVVS
IGTAGDNVTIRFTTDNPGPWFLHCHIDWHLEAGLAVVFAEDTPDIASANPVPAAWDDLCPIYDALSPDD
QPRLQVDLDFQKASSIVYKKEGEQVEFSFPLAFTVEKLTGSGELWWQAERASSSKSWITFDLKNKEVS
VKRVTQDPKLQMGKKLPLHLTLPQALPQYAGSGNLTLALEAKTGKLHQEVNLVVMRATQLQKNLTCEV
WGPTSPKLMLSLKLENKEAKVSKREKAVWVLNPEAGMWQCLLSDSGQVLLESNIKVLPTWSTPVQPM
ALIVLGGVAGLLLFIGLGIFFCVRCRHRRR

Red: IgK signal peptide, Orange: HA epitope tag, Green: V5 epitope tag, Blue: LaccID, Purple: CD4 transmembrane domain

**HA-V5-LaccID(mut)-CD4 TM (Surface LaccID, with active site mutation)**

METDTLLLWVLLLWVPGSTGDYPYDVPDYAGAQPARSGGGKPIPNPLLGLDSTASAIGPVADLHIANA
DISPDGFTRSAVLAGGTFPGPLITGNKGDNFRINVINQLTDTTMLRSTSIHWHGFFQHGTSWADGPAFV
TQCPIAPGNSFLYNFTVPDQAGTFWYHSHLSTQYCDGLSGPFVVYDPNDPHRSLYDVDDESTVITLAD
WYHTPAPQLAAAPPVPDSTLINGKGRYAGGPTSPLAVINVEQGKRYRFRLISISCDPNFTFSIDGHNMTI
IEADGVNTQPLTVDSIQIFAGQRYSFVLNANQPVGNYWIRANPNIGTTGFAGGINSAILRYVGAPEADPT
TTQTTSTNPVNETNLHPLENPGAPGNPTPGGADVNINLNIGFSSTGQFTINGVSFIPPTVPVLLQILSGA
RTAQDLLPSGSVYTLPPNKVIEISMPGGASAAGGPHPFHLHGHTFDVVRSAGSSTYNYVNPVRRDVVS
IGTAGDNVTIRFTTDNPGPWFLAAAIDWHLEAGLAVVFAEDTPDIASANPVPAAWDDLCPIYDALSPDD
QPRLQVDLDFQKASSIVYKKEGEQVEFSFPLAFTVEKLTGSGELWWQAERASSSKSWITFDLKNKEVS
VKRVTQDPKLQMGKKLPLHLTLPQALPQYAGSGNLTLALEAKTGKLHQEVNLVVMRATQLQKNLTCEV
WGPTSPKLMLSLKLENKEAKVSKREKAVWVLNPEAGMWQCLLSDSGQVLLESNIKVLPTWSTPVQPM
ALIVLGGVAGLLLFIGLGIFFCVRCRHRRR

Red: IgK signal peptide, Orange: HA epitope tag, Green: V5 epitope tag, Blue: LaccID (with HCH to AAA mutation), Purple: CD4 transmembrane domain

**HA-V5-HRP-TM (Surface HRP)**

METDTLLLWVLLLWVPGSTGDYPYDVPDYAGAQPARSGGGKPIPNPLLGLDSTASQLTPTFYDNSCPN
VSNIVRDTIVNELRSDPRIAASILRLHFHDCFVNGCDASILLDNTTSFRTEKDAFGNANSARGFPVIDRMK
AAVESACPRTVSCADLLTIAAQQSVTLAGGPSWRVPLGRRDSLQAFLDLANANLPAPFFTLPQLKDSF
RNVGLNRSSDLVALSGGHTFGKNQCRFIMDRLYNFSNTGLPDPTLNTTYLQTLRGLCPLNGNLSALVD
FDLRTPTIFDNKYYVNLEEQKGLIQSDQELFSSPNATDTIPLVRSFANSTQTFFNAFVEAMDRMGNITPL
TGTQGQIRLNCRVVNSNSPRLQVDLDFQKASSIVYKKEGEQVEFSFPLAFTVEKLTGSGELWWQAERA
SSSKSWITFDLKNKEVSVKRVTQDPKLQMGKKLPLHLTLPQALPQYAGSGNLTLALEAKTGKLHQEVN
LVVMRATQLQKNLTCEVWGPTSPKLMLSLKLENKEAKVSKREKAVWVLNPEAGMWQCLLSDSGQVLL
ESNIKVLPTWSTPVQPMALIVLGGVAGLLLFIGLGIFFCVRCRHRRR

Red: IgK signal peptide, Orange: HA epitope tag, Green: V5 epitope tag, Blue: HRP, Purple: CD4 transmembrane domain

**HA-V5-APEX2-TM (Surface APEX2)**

METDTLLLWVLLLWVPGSTGDYPYDVPDYAGAQPARSGGGKPIPNPLLGLDSTASGKSYPTVSADYQ
DAVEKAKKKLRGFIAEKRCAPLMLRLAFHSAGTFDKGTKTGGPFGTIKHPAELAHSANNGLDIAVRLLE
PLKAEFPILSYADFYQLAGVVAVEVTGGPKVPFHPGREDKPEPPPEGRLPDPTKGSDHLRDVFGKAMG
LTDQDIVALSGGHTIGAAHKERSGFEGPWTSNPLIFDNSYFTELLSGEKEGLLQLPSDKALLSDPVFRPL
VDKYAADEDAFFADYAEAHQKLSELGFADAPRLQVDLDFQKASSIVYKKEGEQVEFSFPLAFTVEKLTG
SGELWWQAERASSSKSWITFDLKNKEVSVKRVTQDPKLQMGKKLPLHLTLPQALPQYAGSGNLTLAL
EAKTGKLHQEVNLVVMRATQLQKNLTCEVWGPTSPKLMLSLKLENKEAKVSKREKAVWVLNPEAGM
WQCLLSDSGQVLLESNIKVLPTWSTPVQPMALIVLGGVAGLLLFIGLGIFFCVRCRHRRR

Red: IgK signal peptide, Orange: HA epitope tag, Green: V5 epitope tag, Blue: APEX2, Purple: CD4 transmembrane domain

**HA-V5-LaccID-KDEL (ER LaccID)**
METDTLLLWVLLLWVPGSTGDYPYDVPDYAGAQPARSGGGKPIPNPLLGLDSTASAIGPVADLHIANA
DISPDGFTRSAVLAGGTFPGPLITGNKGDNFRINVINQLTDTTMLRSTSIHWHGFFQHGTSWADGPAFV
TQCPIAPGNSFLYNFTVPDQAGTFWYHSHLSTQYCDGLSGPFVVYDPNDPHRSLYDVDDESTVITLAD
WYHTPAPQLAAAPPVPDSTLINGKGRYAGGPTSPLAVINVEQGKRYRFRLISISCDPNFTFSIDGHNMTI
IEADGVNTQPLTVDSIQIFAGQRYSFVLNANQPVGNYWIRANPNIGTTGFAGGINSAILRYVGAPEADPT
TTQTTSTNPVNETNLHPLENPGAPGNPTPGGADVNINLNIGFSSTGQFTINGVSFIPPTVPVLLQILSGA
RTAQDLLPSGSVYTLPPNKVIEISMPGGASAAGGPHPFHLHGHTFDVVRSAGSSTYNYVNPVRRDVVS
IGTAGDNVTIRFTTDNPGPWFLHCHIDWHLEAGLAVVFAEDTPDIASANPVPAAWDDLCPIYDALSPDD
QKDEL

Red: IgK signal peptide, Orange: HA epitope tag, Green: V5 epitope tag, Blue: LaccID

**HA-V5-HRP-KDEL (ER HRP)**
METDTLLLWVLLLWVPGSTGDYPYDVPDYAGAQPARSGGGKPIPNPLLGLDSTASQLTPTFYDNSCPN
VSNIVRDTIVNELRSDPRIAASILRLHFHDCFVNGCDASILLDNTTSFRTEKDAFGNANSARGFPVIDRMK
AAVESACPRTVSCADLLTIAAQQSVTLAGGPSWRVPLGRRDSLQAFLDLANANLPAPFFTLPQLKDSF
RNVGLNRSSDLVALSGGHTFGKNQCRFIMDRLYNFSNTGLPDPTLNTTYLQTLRGLCPLNGNLSALVD
FDLRTPTIFDNKYYVNLEEQKGLIQSDQELFSSPNATDTIPLVRSFANSTQTFFNAFVEAMDRMGNITPL
TGTQGQIRLNCRVVNSNSKDEL

Red: IgK signal peptide, Orange: HA epitope tag, Green: V5 epitope tag, Blue: HRP

**HA-V5-APEX2-ER (ER APEX2)**
METDTLLLWVLLLWVPGSTGDYPYDVPDYAGAQPARSGGGKPIPNPLLGLDSTASGKSYPTVSADYQ
DAVEKAKKKLRGFIAEKRCAPLMLRLAFHSAGTFDKGTKTGGPFGTIKHPAELAHSANNGLDIAVRLLE
PLKAEFPILSYADFYQLAGVVAVEVTGGPKVPFHPGREDKPEPPPEGRLPDPTKGSDHLRDVFGKAMG
LTDQDIVALSGGHTIGAAHKERSGFEGPWTSNPLIFDNSYFTELLSGEKEGLLQLPSDKALLSDPVFRPL
VDKYAADEDAFFADYAEAHQKLSELGFADAKDEL

Red: IgK signal peptide, Orange: HA epitope tag, Green: V5 epitope tag, Blue: APEX2

**ManII-LaccID-V5 (Golgi LaccID)**
MKLSRQFTVFGSAIFCVVIFSLYLMLDRGHLDYPRGPRQEGSFPQGQLSILQEKIDHLERLLAENNEIISN
IRDSVINLSESVEDGPRGSPGNASQGTRAIGPVADLHIANADISPDGFTRSAVLAGGTFPGPLITGNKGD
NFRINVINQLTDTTMLRSTSIHWHGFFQHGTSWADGPAFVTQCPIAPGNSFLYNFTVPDQAGTFWYHS
HLSTQYCDGLSGPFVVYDPNDPHRSLYDVDDESTVITLADWYHTPAPQLAAAPPVPDSTLINGKGRYA
GGPTSPLAVINVEQGKRYRFRLISISCDPNFTFSIDGHNMTIIEADGVNTQPLTVDSIQIFAGQRYSFVLN
ANQPVGNYWIRANPNIGTTGFAGGINSAILRYVGAPEADPTTTQTTSTNPVNETNLHPLENPGAPGNPT
PGGADVNINLNIGFSSTGQFTINGVSFIPPTVPVLLQILSGARTAQDLLPSGSVYTLPPNKVIEISMPGGA
SAAGGPHPFHLHGHTFDVVRSAGSSTYNYVNPVRRDVVSIGTAGDNVTIRFTTDNPGPWFLHCHIDW
HLEAGLAVVFAEDTPDIASANPVPAAWDDLCPIYDALSPDDQGKPIPNPLLGLDST

Red: ManII, Orange: LaccID, Green: V5 epitope tag

**Aga2p-myc-LaccID (LaccID yeast display)**

MQLLRCFSIFSVIASVLAQELTTICEQIPSPTLESTPYSLSTTTILANGKAMQGVFEYYKSVTFVSNCGSH
PSTTSKGSPINTQYVFKDNSSTIEGRYPYDVPDYAGKPIPNPLLGLDSTENLYFQSLQASGGGGSGGG
GSGGGGSASHMAIGPVADLHIANADISPDGFTRSAVLAGGTFPGPLITGNKGDNFRINVINQLTDTTML
RSTSIHWHGFFQHGTSWADGPAFVTQCPIAPGNSFLYNFTVPDQAGTFWYHSHLSTQYCDGLSGPFV
VYDPNDPHRSLYDVDDESTVITLADWYHTPAPQLAAAPPVPDSTLINGKGRYAGGPTSPLAVINVEQG
KRYRFRLISISCDPNFTFSIDGHNMTIIEADGVNTQPLTVDSIQIFAGQRYSFVLNANQPVGNYWIRANPN
IGTTGFAGGINSAILRYVGAPEADPTTTQTTSTNPVNETNLHPLENPGAPGNPTPGGADVNINLNIGFSS
TGQFTINGVSFIPPTVPVLLQILSGARTAQDLLPSGSVYTLPPNKVIEISMPGGASAAGGPHPFHLHGHT
FDVVRSAGSSTYNYVNPVRRDVVSIGTAGDNVTIRFTTDNPGPWFLHCHIDWHLEAGLAVVFAEDTPDI
ASANPVPAAWDDLCPIYDALSPDDQGSEQKLISEEDL

Red: Aga2p, Orange: HA epitope tag, Green: V5 epitope tag, Blue: LaccID, Purple: Myc epitope tag

**LaccID-8xHis (recombinant LaccID expression)**
MRFPSIFTDVLFAASSALAAPVNTTTEDETAQIPAEAVIGYSDLEGDSDVAVLPFSNGTNNRLLFINTTIA
SIAAKEEGVSLEKRGAEAEFAIGPVADLHIANADISPDGFTRSAVLAGGTFPGPLITGNKGDNFRINVINQ
LTDTTMLRSTSIHWHGFFQHGTSWADGPAFVTQCPIAPGNSFLYNFTVPDQAGTFWYHSHLSTQYCD
GLSGPFVVYDPNDPHRSLYDVDDESTVITLADWYHTPAPQLAAAPPVPDSTLINGKGRYAGGPTSPLA
VINVEQGKRYRFRLISISCDPNFTFSIDGHNMTIIEADGVNTQPLTVDSIQIFAGQRYSFVLNANQPVGNY
WIRANPNIGTTGFAGGINSAILRYVGAPEADPTTTQTTSTNPVNETNLHPLENPGAPGNPTPGGADVNI
NLNIGFSSTGQFTINGVSFIPPTVPVLLQILSGARTAQDLLPSGSVYTLPPNKVIEISMPGGASAAGGPHP
FHLHGHTFDVVRSAGSSTYNYVNPVRRDVVSIGTAGDNVTIRFTTDNPGPWFLHCHIDWHLEAGLAVV
FAEDTPDIASANPVPAAWDDLCPIYDALSPDDQAAALEVLFQGPGSGHHHHHHHH

Red: Signal peptide, Blue: LaccID, Green: HRV 3C protease cleavage site, Purple: 8xHis tag

**HA-V5-LacAnc100-NES (LacAnc100 expression in cytosol)**
MYPYDVPDYAGAQPARSGGGKPIPNPLLGLDSTASAIGPVADLHIVNADISPDGFTRSAVLAGGTFPGP
LITGNKGDNFQINVINQLTDTTMLRSTSIHWHGFFQHGTNWADGPAFVTQCPIAPGNSFLYNFTVPDQA
GTFWYHSHLSTQYCDGLRGPFVIYDPNDPHRSLYDVDDESTVITLADWYHTPAPQLAAAPPVPDSTLIN
GKGRYAGGPTSPLAVINVEQGKRYRFRLISISCDPNFTFSIDGHNMTIIEADGVNTQPLTVDSIQIFAGQR
YSFVLNANQPVDNYWIRANPNIGTTGFAGGINSAILRYVGAPEADPTTTQTTSTNPLNETNLHPLENPG
APGNPTPGGADVNINLNIGFSSTGQFTINGVSFIPPTVPVLLQILSGARTAQDLLPSGSVYTLPPNKVIEIS
MPGGASAAGGPHPFHLHGHTFDVVRSAGSSTYNYVNPVRRDVVSIGTAGDNVTIRFTTDNPGPWFLH
CHIDWHLEAGLAVVFAEDTPDIASANPVPAAWDDLCPIYDALSPDDQASLQLPPLERLTLD

Orange: HA epitope tag, Green: V5 epitope tag, Blue: LacAnc100, Red: Nuclear Export Signal (NES)

**MTS-HA-V5-LacAnc100 (LacAnc100 expression in mitochondria)**
MLATRVFSLVGKRAISTSVCVRAHKDPYPYDVPDYAGAQPARSGGGKPIPNPLLGLDSTASAIGPVADL
HIVNADISPDGFTRSAVLAGGTFPGPLITGNKGDNFQINVINQLTDTTMLRSTSIHWHGFFQHGTNWAD
GPAFVTQCPIAPGNSFLYNFTVPDQAGTFWYHSHLSTQYCDGLRGPFVIYDPNDPHRSLYDVDDESTV
ITLADWYHTPAPQLAAAPPVPDSTLINGKGRYAGGPTSPLAVINVEQGKRYRFRLISISCDPNFTFSIDG
HNMTIIEADGVNTQPLTVDSIQIFAGQRYSFVLNANQPVDNYWIRANPNIGTTGFAGGINSAILRYVGAP
EADPTTTQTTSTNPLNETNLHPLENPGAPGNPTPGGADVNINLNIGFSSTGQFTINGVSFIPPTVPVLLQI
LSGARTAQDLLPSGSVYTLPPNKVIEISMPGGASAAGGPHPFHLHGHTFDVVRSAGSSTYNYVNPVRR
DVVSIGTAGDNVTIRFTTDNPGPWFLHCHIDWHLEAGLAVVFAEDTPDIASANPVPAAWDDLCPIYDAL
SPDDQ

Red: Mito-targeting sequence (MTS), Orange: HA epitope tag, Green: V5 epitope tag, Blue: LacAnc100

**HA-V5-LacAnc100(6 x N-to-A)-TM (LacAnc100 with N-glycosylation mutations)**
METDTLLLWVLLLWVPGSTGDYPYDVPDYAGAQPARSGGGKPIPNPLLGLDSTASAIGPVADLHIVNA
DISPDGFTRSAVLAGGTFPGPLITGNKGDNFQINVINQLTDTTMLRSTSIHWHGFFQHGTNWADGPAFV
TQCPIAPGNSFLYAFTVPDQAGTFWYHSHLSTQYCDGLRGPFVIYDPNDPHRSLYDVDDESTVITLAD
WYHTPAPQLAAAPPVPDSTLINGKGRYAGGPTSPLAVINVEQGKRYRFRLISISCDPAFTFSIDGHAMTII
EADGVNTQPLTVDSIQIFAGQRYSFVLNANQPVDNYWIRANPNIGTTGFAGGINSAILRYVGAPEADPTT
TQTTSTNPLAETNLHPLENPGAPGAPTPGGADVNINLNIGFSSTGQFTINGVSFIPPTVPVLLQILSGART
AQDLLPSGSVYTLPPNKVIEISMPGGASAAGGPHPFHLHGHTFDVVRSAGSSTYNYVNPVRRDVVSIG
TAGDAVTIRFTTDNPGPWFLHCHIDWHLEAGLAVVFAEDTPDIASANPVPAAWDDLCPIYDALSPDDQP
RLQVDLDFQKASSIVYKKEGEQVEFSFPLAFTVEKLTGSGELWWQAERASSSKSWITFDLKNKEVSVK
RVTQDPKLQMGKKLPLHLTLPQALPQYAGSGNLTLALEAKTGKLHQEVNLVVMRATQLQKNLTCEVW
GPTSPKLMLSLKLENKEAKVSKREKAVWVLNPEAGMWQCLLSDSGQVLLESNIKVLPTWSTPVQPMA
LIVLGGVAGLLLFIGLGIFFCVRCRHRRR

Red: IgK signal peptide, Orange: HA epitope tag, Green: V5 epitope tag, Blue: LacAnc100 (Asn at N-X-S/T motifs are mutated to Ala), Purple: CD4 transmembrane domain

**HA-V5-LaccID-CD2 (Fly surface LaccID)**
MDISYIFVICLMALCSGGSSLSQVEGKQKSGRGRGSMWYPYDVPDYAGAQPARSGGGKPIPNPLLGL
DSTASAIGPVADLHIANADISPDGFTRSAVLAGGTFPGPLITGNKGDNFRINVINQLTDTTMLRSTSIHWH
GFFQHGTSWADGPAFVTQCPIAPGNSFLYNFTVPDQAGTFWYHSHLSTQYCDGLSGPFVVYDPNDP
HRSLYDVDDESTVITLADWYHTPAPQLAAAPPVPDSTLINGKGRYAGGPTSPLAVINVEQGKRYRFRLI
SISCDPNFTFSIDGHNMTIIEADGVNTQPLTVDSIQIFAGQRYSFVLNANQPVGNYWIRANPNIGTTGFA
GGINSAILRYVGAPEADPTTTQTTSTNPVNETNLHPLENPGAPGNPTPGGADVNINLNIGFSSTGQFTIN
GVSFIPPTVPVLLQILSGARTAQDLLPSGSVYTLPPNKVIEISMPGGASAAGGPHPFHLHGHTFDVVRSA
GSSTYNYVNPVRRDVVSIGTAGDNVTIRFTTDNPGPWFLHCHIDWHLEAGLAVVFAEDTPDIASANPVP
AAWDDLCPIYDALSPDDQSGRKEITNALETWGALGQDINLDIPSFQMSDDIDDIKWEKTSDKKKIAQFR
KEKETFKEKDTYKLFKNGTLKIKHLKTDDQDIYKVSIYDTKGKNVLEKIFDLKIQERVSKPKISWTCINTTL
TCEVMNGTDPELNLYQDGKHLKLSQRVITHKWTTSLSAKFKCTAGNKVSKESSVEPVSCPEKGLDIYLII
GICGGGSLLMVFVALLVFYITKRKKRRRIPAAAAAAAAV

Red: Signal peptide, Orange: HA epitope tag, Green: V5 epitope tag, Blue: LaccID, Purple: CD2 transmembrane domain
